# Direct prediction of intermolecular interactions driven by disordered regions

**DOI:** 10.1101/2024.06.03.597104

**Authors:** Garrett M. Ginell, Ryan. J Emenecker, Jeffrey M. Lotthammer, Emery T. Usher, Alex S. Holehouse

**Affiliations:** Department of Biochemistry and Molecular Biophysics, Washington University School of Medicine, St. Louis, MO; Center for Biomolecular Condensates (CBC), Washington University in St. Louis, St. Louis, MO

## Abstract

Intrinsically disordered regions (IDRs) are critical for a wide variety of cellular functions, many of which involve interactions with partner proteins. Molecular recognition is typically considered through the lens of sequence-specific binding events. However, a growing body of work has shown that IDRs often interact with partners in a manner that does not depend on the precise order of the amino acid order, instead driven by complementary chemical interactions leading to disordered bound-state complexes. Despite this emerging paradigm, we lack tools to describe, quantify, predict, and interpret these types of structurally heterogeneous interactions from the underlying amino acid sequences. Here, we repurpose the chemical physics developed originally for molecular simulations to develop an approach for predicting intermolecular interactions between IDRs and partner proteins. Our approach enables the direct prediction of phase diagrams, the identification of chemically-specific interaction hotspots on IDRs, and a route to develop and test mechanistic hypotheses regarding IDR function in the context of molecular recognition. We use our approach to examine a range of systems and questions to highlight its versatility and applicability.

## INTRODUCTION

Intrinsically disordered proteins and protein regions (IDRs) are prevalent across the kingdoms of life (*1–5*). While folded domains exist in a stable 3D structure, IDRs exist in a heterogeneous ensemble of conformations. Despite lacking a fixed tertiary structure, disordered regions can play essential roles across many distinct cellular processes (*2*, *5*, *6*). IDRs are frequently involved in molecular recognition, mediating networks of intermolecular interactions that facilitate signaling networks, transcriptional regulation, and translational control. Consequently, there is particular interest in characterizing protein-protein interactions in which one (or both) of the interacting domains are disordered.

IDRs can interact with partner proteins in a variety of ways. Some IDRs (or subregions within IDRs) may fold upon binding, potentially leading to stable bound-state complexes amenable to structural characterization (**Fig. 1A**, *left*)(*7*). These interactions are typically driven by sequence-specific motifs, meaning they depend on the precise order of the amino acids, akin to a conventional structured interface (*8*, *9*). In many cases, however, IDRs do not acquire a stable structure upon binding and exist as a disordered bound-state complex with a folded partner, an interaction mode known as fuzzy binding (**Fig. 1A**, middle) (*10–13*). These more structurally heterogeneous interactions are often driven at least in part by chemical specificity, that is, complementary chemistry between the IDR and its partner (*14*, *15*). Unlike sequence-specific interactions, chemically-specific interactions can tolerate changes to the underlying sequence if chemical complementarity is retained(*14*). Finally, recent work has shown that IDRs can bind other IDRs (or unstructured nucleic acids), and both remain fully disordered in the bound state (**Fig. 1A**, right) (*16–18*). Beyond stoichiometric interactions, the role of IDRs in biomolecular condensates (non-stoichiometric assemblies) has also been extensively investigated. Here, chemically specific multivalent interactions have revealed a rich molecular grammar of IDR-mediated interactions (*19–33*). In short, while investigations of IDR-mediated interactions have historically focussed on sequence-specific binding, a growing body of work suggests chemically-specific interactions may be equally important in tuning or even defining IDR-mediated interactions (*2*, *15*, *34–38*).

**Figure 1.**
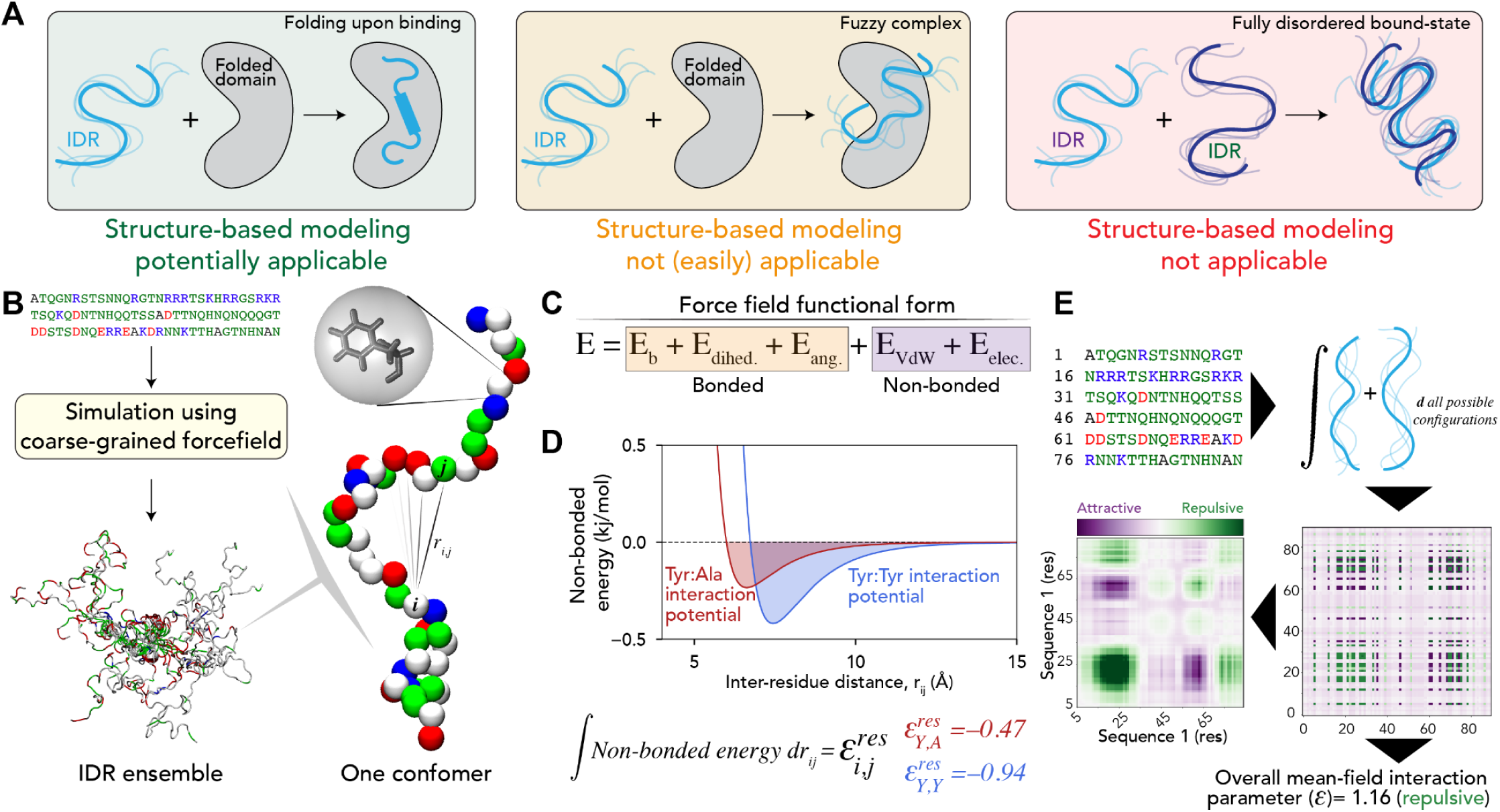
**(A)** IDR intermolecular interaction can occur through a conventional structured interface (left), a disordered bound-state complex with a folded partner (middle), or a disordered bound-state complex where both partners are disordered (right). **(B)** Coarse-grained force fields can represent amino acids as individual beads and generate IDR conformational ensembles by sampling energetically accessible conformers in 3D space. **(C)** Force fields describe bonded and non-bonded components, where non-bonded reflects short-range and long-range interactions that determine attraction or repulsion between individual residues. **(D)** Non-bonded interactions are defined by distance-dependent potentials that describe the relationship between inter-bead distance and instantaneous potential energy. Integrating under these potentials yields a parameter proportional to the overall attraction/repulsion between those beads. **(E)** By assuming two proteins can interact via all possible configurations without concern for chain connectivity, we can calculate inter-residue preferential interaction coefficients and then use local sequence context to convert these into smoothed predicted intermolecular interaction maps (intermaps) or a single mean-filed intermolecular interaction parameter (epsilon, ɛ).

Despite their importance and prevalence, characterizing IDR-mediated interactions—both experimentally and computationally—remains a major challenge. In particular, despite major progress in protein structure prediction, we generally lack ways to predict chemical specificity from sequence alone (*39–41*). Given the emerging importance of chemical specificity in IDR-mediated interactions, our inability to predict these types of recognition hotspots is a major limitation.

Here, we address this knowledge gap through the lens of molecular biophysics. By repurposing the chemical physics developed originally for molecular force fields and discarding requirements for spatial information, we offer a bottom-up framework for predicting chemically-specific IDR-mediated interactions. Our approach is interpretable, and context can be tuned by modulating the underlying physics of the force field. We recognize this approach has many caveats (see *Discussion*), chiefly among them that it is only appropriate for assessing chemical specificity in which an IDR remains largely unstructured. However, given that this is the precise modality to which structure-based approaches are poorly suited, we see our work as complementary to ongoing efforts in the structure prediction space. Predictions are rapid, and our approach is implemented as an open-source Python package (https://github.com/idptools/finches), in Google colab notebooks (https://github.com/idptools/finches-colab), and perhaps most usefully, as an online webserver (http://finches-online.com/) (**Fig. S1**). In short, our work opens the door to high-throughput, straightforward, and interpretable prediction of IDR-mediated intermolecular interactions to guide experiments, predict phase behavior, identify distinct domains, and aid in the interpretation of experimental results.

## RESULTS

### Repurposing molecular force fields for high-throughout bioinformatic analysis of IDR interactions

Molecular force fields describe the chemical physics of biomolecules through a series of equations and parameters (**Fig. 1B, C**). Recent work on coarse-grained models of disordered proteins has led to several force fields that offer accurate predictions of global IDR dimensions, notably among those of the Mpipi and CALVADOS families (*42–45*). These models prescribe a set of equations and parameters that quantify the nonbonded interactions between every pair of amino acids to describe the attraction or repulsion of a pair of residues at some arbitrary distance (**Fig. 1D**). We reasoned this chemical physics – while generally used for simulation – could be stripped out and repurposed by taking the integral under the pairwise potential as a means to calculate a mean-field interaction parameter between two residues, akin to a Mayer-f function without a volume correction component (see *Supporting Information*) (**Fig. 1D**). Using this force field-derived interaction parameter as a starting point and then tuning the interaction of individual amino acids based on their local context (for charged and aliphatic hydrophobic residues, specifically), the resulting inter-residue matrix between a pair of residues can be averaged to obtain a single mean-field inter-protein interaction parameter epsilon (ɛ), or averaged over a sliding window to decode local intermolecular interactions that are attractive or repulsive (**Fig. 1E, Fig. S1**). We refer to the predicted intermolecular interaction maps as “intermaps,” as shown in the bottom left of **Fig. 1E**. In all cases, negative ɛ values are attractive, and positive ɛ values are repulsive.

A central assumption in this approach is that the attraction between two IDRs is mediated solely by complementary chemical interactions (chemical specificity) that emerge in the limit of all possible configurations being sampled, not via precise “structured” interaction between subregions. It also does not allow coordination between distinct regions, bringing with it a host of caveats and considerations (see *Discussion* & *Supporting Information*). Nevertheless, this approach enables us to easily calculate a mean-field interaction parameter, as well as identify subregions that are expected to drive attractive and repulsive interactions. Importantly, the molecular bases for such interactions are entirely interpretable and codified by the underlying functional form of the force field and its associated parameters.

In this work, we implement this approach using the Mpipi-GG and CALVADOS2 force fields, although additional force fields could easily be implemented (*43*, *45*). These models both allow us to modulate the solution environment to whatever extent a force field is parameterized (e.g., here in terms of salt via a Debye-Hückel term). It also allows high-throughput prediction (>1000 sequences per second for a 100-residue IDR, **Fig. S2**). While we emphasize the resulting interaction scores are not expected to offer high-resolution, quantitative predictions, they enable rapid semi-quantitative descriptions of likely IDR-associated interactions.

### Validation against molecular interaction

As an initial test, we first asked if the mean-field interaction parameter we calculate from sequence (ɛ) is proportional to experimentally-measurable values for intermolecular interaction. In principle, a mean-field interaction parameter should be approximately proportional to the osmotic or light-scattering second virial coefficients (B_2_ and A_2_, respectively) (*46*). B_2_ and A_2_ are experimentally measurable quantities that report on the deviation from so-called “ideal behavior,” where ideality here reflects non-interacting molecules. A negative B_2_ or A2 implies net attractive intermolecular interactions, while a positive B_2_ or A_2_ implies a net repulsive intermolecular interaction. We considered two systems where second virial coefficients have previously been characterized for IDRs: variants of the low-complexity domain of the RNA binding protein FUS and the RGG domain from the DEAD-box helicase LAF-1 (*47*, *48*). For FUS, A_2_ values were calculated for a series of mutants (**Fig. 2A**), with values correlating well with predicted ɛ values (see **Fig. S3**). Despite distinct scales, the value of 0 should be equivalent in ɛ and A_2_ space, a prediction confirmed by the fact the best-fit line travels through 0,0 (**Fig. 2A**). Intermaps comparing wildtype FUS with the tyrosine-to-serine (Y2S) mutant illustrates the complete suppression of attractive interactions (**Fig. 2B**). For the RGG domain, we calculated the salt-dependent ɛ values and compared them against NaCl-dependent B_2_ values, yielding a 1:1 correspondence between measured values and predictions (**Fig. 2C**). While there are many caveats associated with relating second virial coefficients to mean-field interaction parameters (see *Supplementary Information*), the trends here gave us confidence that our underlying assumptions were reasonable.

**Figure 1.**
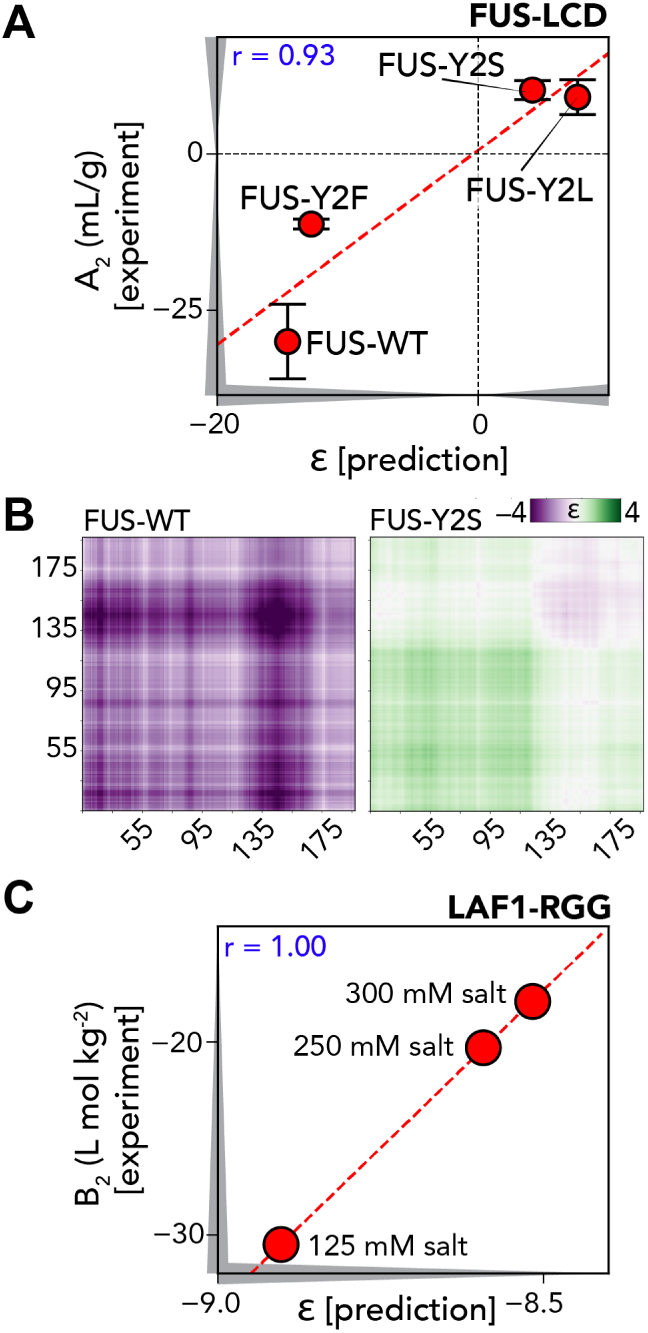
**(A)** Comparison of the light-scattering second virial coefficient (A_2_, which under dilute conditions is sirongly correlated with the osmotic second virial coefficient) with overall ∈ value for different IDRs (data taken from Lin *et al*. 2018). Note experiments are done with FUS and its variants tethered to a common scaffold system (see *Supporting Information*). **(B)** Intermaps for FUS and FUS-Y2S, illustrating how inter-molecular interactions vary. (C) Comparison of the osmolic second virial coefficient with overall ∈ value for the same IOR under different solution conditions. Here, salt weakens iniermolecular interactions (∈ becomes less negative).

### Direct prediction of phase diagrams from sequence

Recent interest in biomolecular phase separation has led to a number of experimental studies characterizing full-phase diagrams of disordered proteins *in vitro*. While predicting phase behavior from sequence has been a goal for many predictors, conceptual and technical challenges have limited their generalizability and scope (see Supplementary Information). With this in mind, we investigated whether our mean-field approximation could recapitulate known relationships between IDR sequence and phase behavior.

We first sought to use ɛ values as input to Flory-Huggins theory, from which phase diagrams can be predicted. Flory-Huggins theory is a simple mean-field solution mixing theory that considers the balance between entropy and enthalpy to determine if a solution of a given temperature and composition will exist in a single phase or multiple phases (*49–51*). Although Flory-Huggins theory is reductive, our goal is not to quantitatively reproduce co-existence curves to match 1:1 with experiments but to provide qualitative predictions for how changes in environment or sequence are expected to alter phase diagrams. To achieve this, we calculated homotypic ɛ values for proteins where full phase diagrams have previously been measured, converted the (extensive) ɛ into an (intensive) Flory χ parameter by dividing by the sequence length, and used the recently developed analytical solution to the Flory-Huggins model of Qian *et al*. to solve full phase diagrams for a series of systems (**Fig. 2A**) (*52*). For a detailed overview of how to read phase diagrams, see **Fig. S4A**. Our predicted phase diagrams report temperature normalized by the critical temperature of a reference sequence (T/T_C_) and concentration as volume fraction (Φ). Phase diagrams here were predicted with the Mpipi-GG-based ɛ analysis, but equivalent results are obtained using CALVADOS2-based analysis.

We began with previously characterized aromatic variants of the low-complexity domain of the RNA binding protein hnRNPA1, a 135-residue low-complexity prion-like domain (**Fig. 3B**) (*20*). These variants increase (Aro^+^) or decrease (Aro^-^, Aro^--^) the number of aromatic residues in the sequence. Our approach yielded full-phase diagrams that show good agreement with respect to the relative impact of aromatic mutations (**Fig. 3C**). Prior work to elucidate these phase diagrams combined simulations and experiments to arrive at conclusions regarding the impact of aromatic residues on phase behavior. In contrast, the major benefit of our approach is these phase diagrams can now be predicted directly from the sequence in seconds.

**Figure 3.**
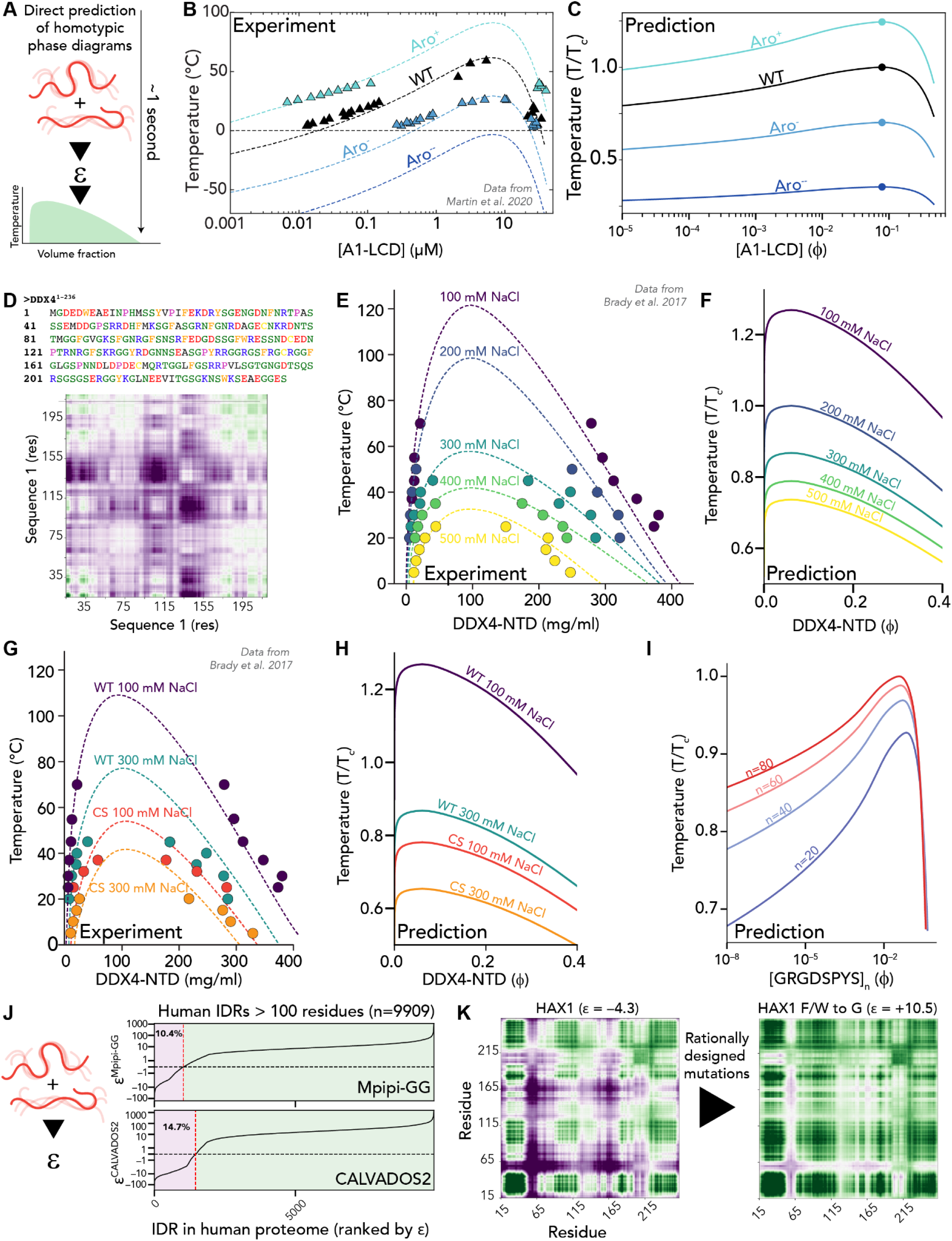
**(A)** Using ɛ as a means to parameterize a Flory-Huggins description of phase behavior, our ɛ-based approach enables the prediction of sequence- and solution-dependent full phase diagrams in seconds. **(B)** Experimentally measured phase diagrams for four variants of the low-complexity domain (LCD) from the RNA binding protein hnRNPA1. These measurements were originally reported by Martin et al. and are reproduced here for easy comparison with predictions (*20*). **(C)** Direct predictions of full phase diagrams using only sequence as input. **(D)** The N-terminal IDR from the DEAD-box helicase protein DDX4 (DDX4-NTD) is a polyampholytic disordered protein that undergoes phase separation *in vitro*. Intermaps predict both clusters of charged residues and aromatic residues contribute orthogonal chemical interactions to drive phase separation. **(E)** Experimentally measured phase diagrams for wildtype DDX4-NTD under four different salt concentrations. These measurements were originally reported by Brady et al. and are reproduced here for easy comparison with predictions (*59*). **(F)** Direct predictions of salt-dependent full-phase diagrams using only sequence and salt concentration as input. **(G)** Experimentally measured phase diagrams for wildtype and charge shuffle (CS) DDX4-NTD variants under two different salt concentrations. These measurements were originally reported by Brady et al. and are reproduced here for easy comparison with predictions (*59*). **(H)** Direct predictions of sequence and salt-dependent full-phase diagrams using only sequence and salt concentration as input. Note that in all cases, the composition of the sequence is identical here, salt and the order of amino acids are varying. **(I)** Direct predictions of length-dependent phase diagrams for resilin-like polypeptide (RPL) [GRGDSPYS]_n_. These phase diagrams are in good agreement with dilute-phase binodal measured by Dzuricky et al. (*62*)**. (J)** Proteome-wide analysis of homotypic ɛ values for all IDRs in the human proteome above 100 amino acids. **(K)** Example analysis comparing overall ɛ and intermaps for the disordered protein HAX1, illustrating how intermaps enable rational mutagenesis to re-wire predicted intermolecular interaction properties.

We next asked how well our approach could capture the solution environment and sequence patterning. Sequence patterning refers to the relative positions of amino acids along the sequence, where the patterning of charged residues, in particular, has been shown to modulate phase behavior in various systems (*53–58*). The N-terminal intrinsically disordered domain of the RNA helicase DDX4 (DDX4-NTD) has been extensively studied in this context (**Fig. 3D**) (*21*, *59–61*). Intermaps identify both charge clusters and aromatic residues as key drivers of intermolecular interaction (**Fig. 3D, bottom**). Prior work measured full phase diagrams as a function of NaCl (**Fig. 3E**), which are correctly reproduced with our approach (**Fig. 3F**). Moreover, Charge Shuffle (CS) variants that maintain the same composition but reposition a small number of charged residues lead to changes in the phase diagram that are correctly recapitulated by our approach, as are arginine-to-lysine and phenylalanine to alanine mutants that entirely suppress phase behavior (**Fig. 3G, 3H, Fig. S4).** Taken together, these results illustrate that our approach is capable of capturing effects driven by sequence patterning, changes in the identity of cationic residues, and the solution environment.

We also assessed how chain length tunes the phase behavior of a resilin-like polypeptide (RLP) construct. As expected, longer chains shift the critical temperature up and the saturation concentration down, recapitulating experimental results (*62*) (**Figure 3I**). Beyond these examples, we predicted full-phase diagrams for a range of systems examined previously, including variants of hnRNPA1-LCD, FUS, and RLP (**Fig. S5**). We also performed an extensive investigation into the low-complexity domain of TDP-43, highlighting that our approach correctly integrates the density of aliphatic residues in the TDP-43 conserved region, recovering experimentally reported consequences of changing a variety of different chemistries (**Fig. S6**) (*63*, *64*). In all cases tested, our predictions (at least qualitatively) capture the effects of sequence chemistry on phase behavior and offer clear explanatory power for how sequence changes are expected to alter intermolecular interactions in the context of IDR-mediated phase separation.

Given the performance of our model, we next calculated homotypic ɛ values for all long IDRs in the human proteome (“long” here is defined here as having over 100 residues) using two different force field-based approaches (Mpipi-GG and CALAVDOS2) (**Supplementary Table S1, S2**). We saw differences in the fraction of IDRs with attractive homotypic ɛ values (i.e., ɛ <0), with ∼10% of IDRs using Mpipi-GG and ∼15% using CALVADOS2 (**Fig. 3J**). These differences reproduce previously reported differences in IDR global dimensions between the two models, whereby Mpipi ensembles were slightly more expanded than CALVADOS-derived ensembles (*45*, *65*). An attractive homotypic ɛ value does not necessarily mean the IDR is predicted to undergo homotypic phase separation in a biochemically relevant context (see **Fig. S7A,B**). However, it does imply the IDR has the potential to self-interact. GO analysis of proteins with IDRs that have attractive homotypic ɛ values (vs. all proteins with long IDRs) identifies RNA-associated processes, morphogenesis, and development as key biological processes associated with these proteins (**Fig. S7C,D**). The enrichment for RNA binding proteins again qualitatively agrees with recent proteome-wide analyses on IDR compaction, highlighting the symmetry between intramolecular interactions and homotypic intermolecular interactions (**Fig. S7**) (*20*, *45*, *65*, *66*). As a final note, we emphasize explicitly and unconditionally that we make no claims whatsoever as to the physiological relevance of these predicted attractive homotypic interactions.

Given that IDR-mediated chemical specificity depends on the amino acid-encoded chemistry, post-translational modifications offer one route to recode that chemistry. To this end, we calculated all homotypic ɛ values for all human IDRs that possess one or more phosphosite (19,703 IDRs) before and after making phosphomimetic mutations. Only experimentally-reported phosphosites (S/T/Y) were used, and in all cases, were converted to E (glutamic acid) (*67*, *68*). Interestingly, ∼57% of IDRs that undergo phosphorylation showed a reduction in homotypic attractive interaction upon phosphorylation (**Fig. S8**), while ∼30% showed an increase in homotypic attractive interaction (**Supplementary Table S3**). Overall, our work reveals many IDRs poised for homotypic (and likely heterotypic) interaction and suggests that their underlying chemical specificity can be rewired through post-translational modifications.

While the overall ɛ value provides a simple mean-field description of the average interprotein interaction, we anticipate the most useful application of these analyses will be in building IDR-centric intermaps. As an example, we highlight the largely disordered protein HAX1, a hub protein involved in cortical actin organization and endocytosis (*69–71*). Intermaps predict phenylalanine and tryptophan residues will drive attractive intermolecular interactions at several specific locations (**Fig. 3J**). Based on this analysis, an F/W to G mutant suppresses those interactions and is predicted to abrogate homotypic intermolecular interaction. We have no reason *a priori* to think HAX1 interacts homotypically, and suggest these aromatic residues would likely drive interaction with other partners, too. However, in the limit of designing targeted and specific mutations to enhance or suppress homotypic (or heterotypic) interaction, our approach offers a clear route to developing testable hypotheses.

### Organizing IDRs in intermolecular chemical space

Given IDRs are often highly variable across different species, there has been substantial interest in understanding if and how one can identify homologous IDRs without using multiple sequence alignments. Our approach opens the door to high-throughput bioinformatics, whereby similarity is considered in terms of chemical space. With this in mind, we wondered how homotypic vs. heterotypic interactions between IDRs varied across the human proteome. We identified all IDRs in the human proteome between 100 and 150 residues in length (3,414 IDRs), focussing on this specific size to ensure intermolecular pairs were approximately equal in length. Next, we computed all possible pairwise interactions (around 12 million calculations), allowing us to map the heterotypic interaction landscape at the proteomic scale (**Fig. 4A**).

**Figure 4.**
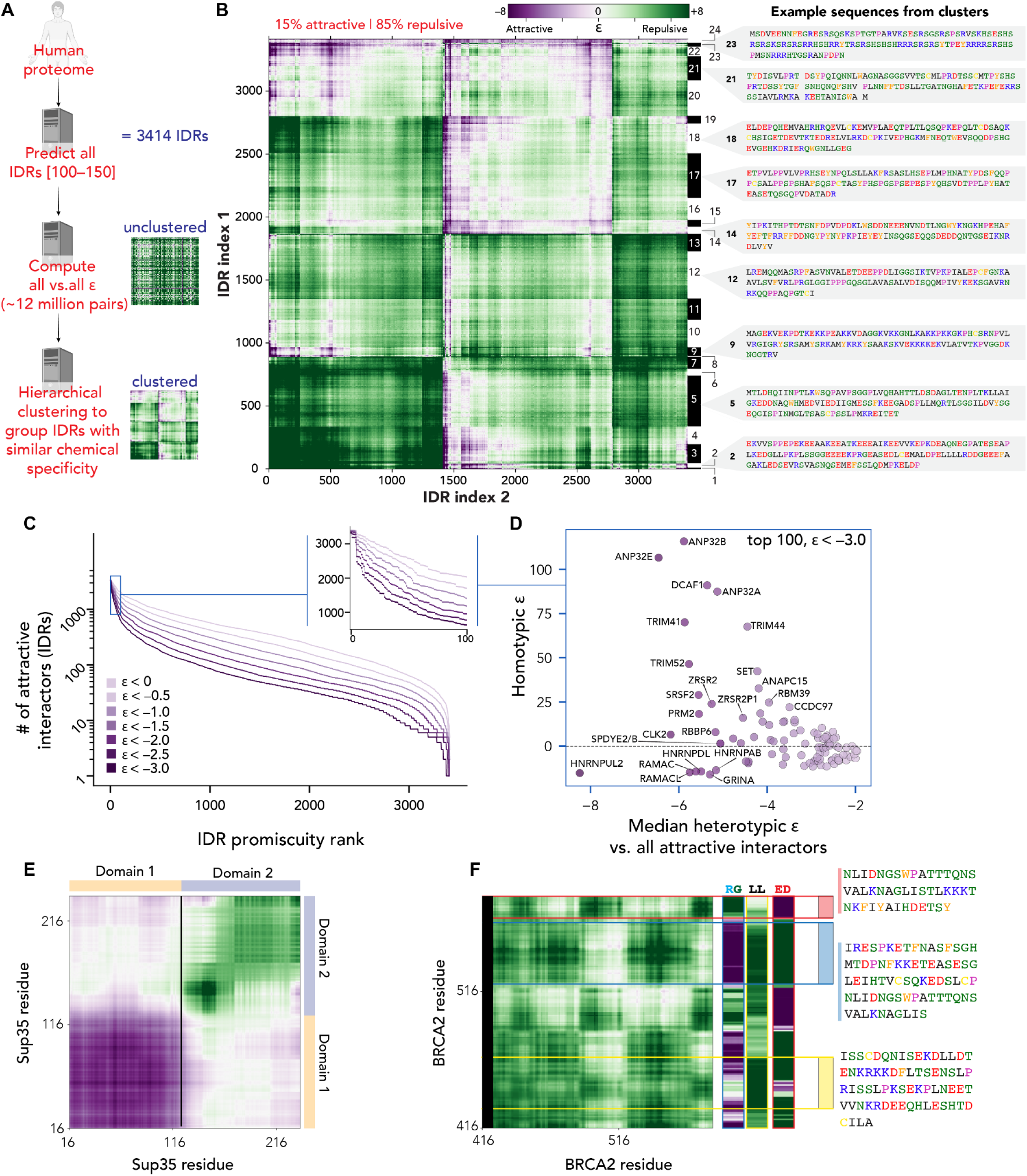
**(A)** Overview of workflow for construction of heterotypic chemical interaction map. **(B)** 3414 IDRs clustered via hierarchical clustering into groups that show similar global intermolecular interaction fingerprints (**Supplementary Table S4**). The average chemical properties of sequences in each cluster are shown in **Supplementary Table S5 (C)** All IDRs ranked by the number of attractive interactions where an ‘attractive interaction’ is defined as ɛ below some threshold value indicated by the purple hue (**Supplementary Table S6**). Inset shows zoom-in on the top 100 proteins. **(D)** Comparison of median heterotypic ɛ vs. homotypic interaction ɛ for the top 100 proteins reveals some highly promiscuous proteins that are also predicted to interact strongly homotypically (homotypic ɛ < 0), while others are predicted to be obligatory heterotypic interactors (homotypic ɛ > 0). Gene names for a subset of the proteins are shown when graphically convenient. **(E)** Domain decomposition of disordered regions in the yeast prion protein Sup35 based on homotypic intermaps. **(F)** Domain decomposition of a BRCA1 subregion using a subset of pre-defined chemically distinct chemical fingerprints.

Hierarchical clustering into 24 clusters revealed subsets of IDRs that showed similar chemical specificity interaction fingerprints, despite being highly diverse in terms of their absolute sequence. Of note, only 15% of heterotypic IDR:IDR interactions are attractive, with the majority being repulsive. While overall repulsive ɛ does not mean two IDRs cannot interact, our work here illustrates two key points: first, when classified in terms of mean-field intermolecular interaction, naturally occurring IDRs fall into distinct chemical niches. Second, sequence chemistry determines whether an IDR is poised for homotypic or heterotypic attractive interactions, and some chemistries appear much more promiscuous than others (*26*).

We next sought to explore the idea of chemical promiscuity further. We define chemical promiscuity as the tendency for an IDR to possess attractive ɛ values for a large number of different potential partners. To quantify this, we ranked each IDR by the number of attractive heterotypic ɛ values (**Fig. 4C**). This analysis reveals many IDRs have the potential to be highly promiscuous. Excising the top 100 most promiscuous IDRs, we identified a range of molecular functions, notably RNA binding, but also proteins involved in cellular homeostasis (e.g., TRIM41, DCAF1, ANAPC15), apoptosis (ANP32B, ANP32A, SET, CLK2, GRINA), transcriptional regulation (ANP32A, SET), and histone chaperoning (ANP32E, SET) (**Supplementary Table S6**). Moreover, taking protein copy number information into account, high-abundance and promiscuous IDR-containing proteins are almost universally RNA-binding proteins (*72*) (**Fig. S9**). In short, while most possible IDR:IDR interactions are repulsive (i.e., most rows in Fig. 4B are largely green), all IDRs do interact favorably with at least one other IDR (i.e., each row in Fig. 4B has at least one purple pixel), and many IDRs are in principle highly promiscuous (i.e., some rows in Fig. 4B are largely purple, e.g., clusters 23, 24).

The ability to segment IDRs based on intermolecular chemical specificity is not limited to proteome-scale analyses. While identifying IDRs in proteins is now relatively straightforward, sub-classification of internally distinct subdomains has been a historically challenging exercise. A major confounding factor here is that IDR function is context-specific; as such, a “functional domain” only makes sense to define in the context of some function. If we restrict ourselves to chemically specific molecular interactions as our function of interest, it becomes possible to define distinct subdomains within an IDR in the context of some interaction partner. For a given pair of IDRs, we can segment subregions into chemically distinct domains, offering clear guidelines for subdomain deletion studies beyond arbitrary cut-off points (**Fig. 3E**). This is illustrated here in proteins with IDRs with clear compositional biases (e.g., the N-terminal half of the yeast prion protein Sup35) but is perhaps most useful for segmenting large IDRs where interaction partners are not yet known using a limited set of chemical fingerprints (**Fig. S10, S11**) (e.g., the highly disordered protein BRCA1).

### Decoding chemical-specificity for IDR-mediated interactions

Finally, to determine how well our approach can help us explain and uncover intermolecular interactions between IDRs in the context of protein-protein interactions, we examined a set of previously studied systems. Our goal is to determine whether intermaps enable the development of testable and well-motivated hypotheses regarding intermolecular interactions between an IDR and an associated partner. By examining previously studied systems, we assess whether observed behavior could have been predicted *a priori*.

Prothymosin alpha (ProTα) and Histone H1 (H1) co-assemble into a fully disordered complex with picomolar affinity (**Fig. 5A**) (*16*). ProTα is entirely disordered, while H1 contains a small globular domain flanked by N and C-terminal disordered regions (**Fig. S12**). While the two proteins assemble via complex coacervation, we wondered if our ɛ-based analysis would allow us to discern specific subregions that contribute more or less to binding. Extant binding data obtained from single-molecule FRET (smFRET) showed that the C-terminal half of H1 binds ProTα with a K_D_ of 0.04 nM, while the N-terminal region binds much less tightly with a K_D_ of 173 nM (**Fig. 5B**). Gratifyingly, by calculating the per-residue sum of ProTα:H1 intermaps (**Fig. S12**), we find a stark difference in the predicted interaction strength between the two halves (**Fig. 5C**). This effect is captured for both Mpipi- and CALAVDOS-based models, with both predicting that the C-terminal half will bind much more strongly than the N-terminal half (**Fig. 5D**). In addition, a strong salt-dependence on this interaction is predicted, in line with published work (**Fig. S12**). Taken together, this illustrates that even for relatively low-complexity, high-affinity interactions, local chemical specificity is, in principle, predictable.

**Figure 5.**
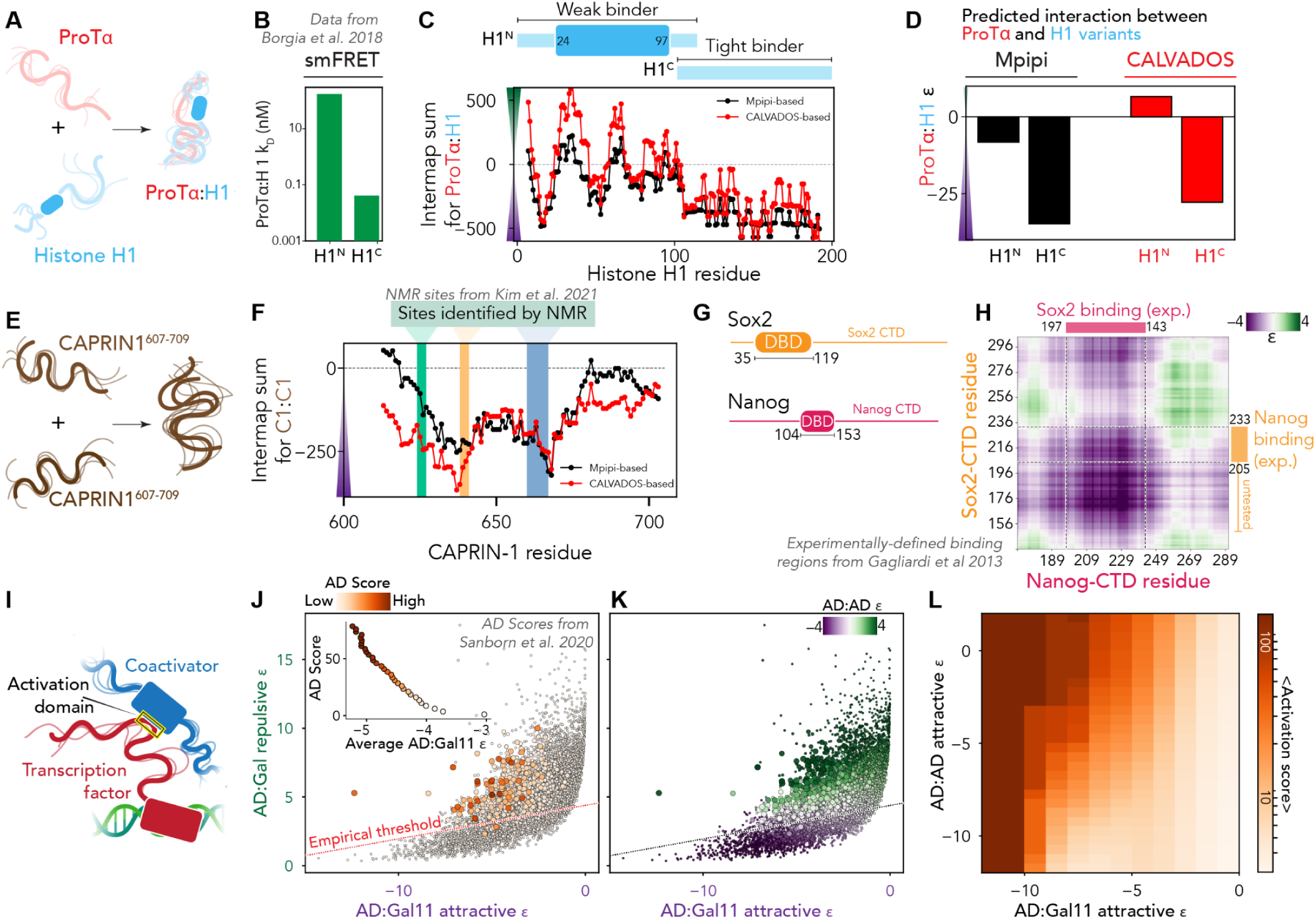
**(A)** Overview of the ProTα and H1 system. **(B)** Previously measured binding data for the N- and C-terminal halves of ProTα. These measurements were originally reported by Borgia *et al.* and are reproduced here for easy comparison with predictions (*16*). **(C)** Per-residue intermap sum for H1 with ProTα, where values come from summing over allow rows in each H1 residue column from the intermap (**Fig. S12**). More negative values mean more attractive. **(D)** Sum of N and C-terminal intermap sums to capture gross relative affinities of the two halves in both Mpipi and CALVADOS. **(E)** Overview of the CAPRIN-1^607-709^ homotypic interaction. **(F)** A comparison of per-residue intermap sum for CAPRIN-1^607-709^ homotypic interaction with NMR hotspots is highlighted. NMR hotspots were originally reported by Kim *et al.* and are reproduced here for easy comparison with predictions (*73*). **(G)** Domain schematic of transcription factors Sox2 and Nanog with DNA binding domains highlighted. The remainder of the proteins are disordered (**Fig. S14**). **(H)** The intermap between Sox2 and Nanog CTDs, with the previously identified binding region highlighted, is Note that Sox2^156-205^ has not been tested as mediating Nanog interaction but is strongly predicted to be critical for this binding. In both cases, the specific residues that are predicted mediate this interaction (Trp in Nanog and Tyr in Sox2, **Fig. S14**) also underlie this interaction experimentally(*74*). **(I)** Overview of our working model for transcription-factor coactivator interaction. **(J)** Scatter plot comparing attractive and repulsive ɛ values for activation domain (AD):(Gal11^158-238^) interaction for all tiles measured by Sanborn et al., where marker size and color reports on activation domain score. The empirical threshold line separates regions where pairs are predicted to have strongly attractive AD:Gal11 interaction, yet no strong activation domain tiles are reported. Inset shows a strong correlation between tiles with a strong AD score and tiles that interact favorably with Gal11^158-238^. **(K)** Same points and sizes as in panel J, with markers colored by homotypic ɛ value. Markers below the empirical threshold show strong homotypic interaction. **(L)** Map of average activation domain activity given some combination of homotypic ɛ (y-axis) and AD:Gal11 ɛ (x-axis). AD:AD (y-axis) sets a threshold where tiles in each pixel have a homotypic ɛ below (more favorable than) the y-axis value. AD:Gal11 (x-axis) sets a threshold where tiles have an attractive ɛ below (more favorable than) the x-axis value. Attractive AD:AD interactions are antagonistic to the average AD score, whereas attractive AD:Gal11 drives the average AD score.

Another example of a dynamic interaction is that of the homotypic interaction between a fragment from the C-terminal IDR of the stress granule-associated protein CAPRIN-1 (**Fig. 5E, Fig. S13**) (*73*). Recent NMR work characterized three interaction hotspots (residues 624–626, 638–640, and 660–666, highlighted as green, yellow, and blue, respectively) that contribute key interactions to CAPRIN-1 intermolecular behavior (**Fig. 5F**). In line with this, two of the three hotspots are clearly predicted from the sequence, with a smaller peak shown for the first. Moreover, unlike DDX4 (for which salt suppresses phase separation), CAPRIN-1 homotypic phase separation is enhanced at higher salt concentrations, an effect also recapitulated by our approach (**Fig. S13**). This result highlights how the complex interplay of charged and aromatic residues can be appropriately captured.

Finally, the master transcription factors Sox2 and Nanog have previously been shown to interact via their C-terminal IDRs (**Fig. 5G**) (*74*). The subregions that drive this interaction are correctly predicted from a heterotypic intermap, highlighting the potential for intermaps to help prioritize the exploration of subregions within disordered domain proteins that may drive intermolecular interactions (**Fig. 5H**). This approach also predicts that the C-terminal IDR of Nanog will undergo robust homotypic phase separation, in agreement with recently published work (**Fig. S14**). This result highlights how - given two proteins of interest - likely chemically-specific sites of intermolecular interaction can be readily predicted.

Having examined several specific examples of IDR-IDR interaction, we asked if IDR:folded domain interactions could also be assessed. Transcription factors are DNA-binding proteins that typically consist of a folded DNA binding domain and an intrinsically disordered region (**Fig. 5I**) (*75*, *76*). Among various functions, transcription factor IDRs can recruit co-activators (e.g., Med15, Gal11 in yeast) which in turn drive transcription(*77–79*). Recent work from several groups has reported detailed high-throughput studies to identify sequence features associated with so-called activation domains (ADs) - sub-regions within transcription factor IDRs that drive gene expression through the recruitment of co-activators (*79–86*).

In yeast-based high-throughput assays, the presumed core co-activator is Gal11. Here, activation domain recognition is performed at least in part by the activation domain binding domain (ABD1) on Gal11 (*87*). We wondered if we could calculate mean-field attractive and repulsive interactions between the surface of the ABD1 and short disordered sequences assayed previously to explain reported activation domain scores (i.e., a measure of how robustly a specific sequence drives gene expression). Using the solvent-accessible residues on the Gal11 ABD1 (PDB 2LPB:A, **Fig. S15**), we calculated IDR:folded domain surface attractive and repulsive values for each of the 7577 40-residue tiles measured previously by Sanborn *et al.* in a high-throughput screen (*79*). Plotting attractive AD:Gal11 interaction vs. repulsive AD:Gal11 for each tile, there was a clear bias for attractive sequences in those that had a higher AD score (**Fig. 5J**). By classing each tile as either a “strong” or “weak” AD, where strong is defined by a variable AD score threshold, we calculated the average AD:Gal11 interaction for strong ADs and found a robust correlation (*r* = 0.94) between average AD score and average attractive interaction (**Fig. 5J**, inset). This conclusion mirrors mRNA display experiments done by Sanborn et al., with the key difference being here we can predict the interaction biases directly from sequence. In summary, these results show that tiles that drive robust gene expression are also generally predicted to interact strongly with Gal11, illustrating the power of this approach for uncovering chemical specificity from high-throughput experiments.

While Gal11 interaction appears to be strongly correlated with AD score, we were surprised to see many tiles with stronger predicted Gal11 interaction yet low AD scores (**Fig. 5J**). A distinct line in the data appeared to dissect the distribution, where almost no tiles under the line had high AD scores (**Fig. 5J, empirical line**). On manual inspection of the tiles under this line, we noticed an abundance of aromatic residues, as well as the occasional arginine or lysine. Calculating the AD:AD ɛ scores, we realized all these sequences were predicted to engage in strong, attractive homotypic interactions (**Fig. 5K**). In effect, our analysis suggested that strong homotypic interaction is detrimental to activation domain function.

To investigate this further, we calculated the average AD scores for sequences with an attractive AD:AD interaction below (more favorable than) some threshold and with an attractive AD:Gal11 interaction below (more favorable than) a second threshold (**Fig. 5L**). Our analysis clearly reveals that the combination of AD:Gal11 and AD:AD interaction determines the AD score of a given sequence. For a given AD:Gal11 strength, AD score can be tuned up or down by suppressing or enhancing AD:AD interaction.

Our work suggests two mechanistic tenets of activation domain function: (i) they must interact strongly with co-activators, and (ii) they must avoid interacting strongly with other partners, here quantified in terms of homotypic interaction. These results are largely consistent with data interpreted by the Acidic Exposure model originally proposed by Staller & Cohen, as well as experimental data from the original study (*79*, *80*). Of note, some activation domains have their activity enhanced upon the addition of hydrophobic aromatic residues, whereas, for others, the addition of aromatic residues has no effect or may even be detrimental (*79*, *81*, *84*, *85*). We suggest our conclusion here offers an explanation for why this may be the case.

While we focus on homotypic interaction out of necessity, we are agnostic as to whether the underlying molecular determinant here is (1) intramolecular interaction (i.e., as proposed by the Acidic Exposure mode), (2) homotypic intermolecular interaction or (3) heterotypic interaction with another cellular component (nucleic acids, proteostatic machinery, the nuclear pore complex, *etc.*). In summary, the analysis highlights how our approach can aid in the analysis and interpretation of large-scale studies using molecular biophysics as a lens for interpretation.

## DISCUSSION & CONCLUSION

By excising the chemical physics developed for molecular simulations, we have repurposed the analytical forms and molecular parameters used to describe inter-amino acid interaction as a means to estimate IDR-associated attractive and repulsive interactions directly from sequence. While many caveats remain, our approach is fast, simple, and offers the ability to perform proteome-scale intermolecular interaction predictions.

Where tested, we have found predictions made by our approach offer qualitative-to-semi-quantitative insight into a wide variety of different systems. This includes phase diagram prediction (**Fig. 3**, **Fig. S4, S5, S6, S8, S13, S14**), classification and domain definition of IDRs based on sequence chemistry (**Fig. 4**) or elucidating sequence-dependent intermolecular interactions between IDRs and their partners (**Fig. 5**). We see this approach as being complementary to existing IDR-associated analyses, offering a means to prototype and develop conceptual hypotheses regarding the likely role(s) of IDRs or subregions in the context of intermolecular interaction.

By computing thousands of heterotypic intermolecular ɛ values, we were able to cluster around 3000 IDRs from the human proteome into chemically similar groups (**Fig. 4**). Some clusters showed strong biases for particular sequence chemistries (**Supplementary Table S5**). However, several large clusters (e.g., clusters 5, 12, 16, and 17, consisting of 38% of all sequences) lacked strong average chemical biases, with a net neutral charge and an intermediate fraction of charged residues but a higher fraction of aliphatic and polar residues. We suggest these more chemically neutral sequences may either be enriched for short linear motifs or may have multiple chemically distinct subdomains, averaging out to a more neutral mean behavior. Further work to explore the substructure of these clusters is ongoing, as well as delineating individual IDRs into chemically distinct subdomains (see below).

One possible application of our approach is in the prediction of homotypic phase diagrams directly from sequence (**Fig. 3**). While one could envisage applying our approach to interrogate the underlying molecular grammar associated with phase separation across disordered regions, prior and ongoing work from other groups has exhaustively explored the underlying chemical principles encoded by coarse-grained forcefields (*24*, *42*, *44*, *88–95*). We build on this prior work, moving away from the need to extrapolate general principles to specific systems and instead enable direct predictions for how mutations are predicted to impact phase diagrams – at least qualitatively – on a case-by-case basis.

Recent work by von Bulow *et al.* has taken an innovative approach combining active learning with coarse-grained simulations to generate the requisite data to train a machine learning model to predict homotypic phase diagram properties (i.e. concentrations in the dense and dilute phase) from sequence (*95*). We suggest if quantitative insight into *in vitro* saturation concentrations is desired, this approach is likely much more robust than ours.

While much of the comparison done here in this manuscript is in the context of phase separation, looking forward, we anticipate that our approach will be most useful in three distinct areas unrelated to biomolecular phase separation.

First, we see the most immediate impact of our approach in the guidance and interpretation of experiments examining IDR-mediated intermolecular interactions. We and others subscribe to an emerging model whereby IDR-mediated interactions are driven by a combination of sequence-specific and chemical-specific interactions (*2*, *14*, *15*, *34*). Our approach provides a means for anyone to easily and quickly quantify chemical specificity between an IDR and a partner. Importantly, because intermaps offer chemically specific insight into how an IDR may interact, they are effectively instantaneous hypothesis generators with respect to understanding the mapping between IDR sequence and molecular interaction. Beyond binary interactions, we anticipate our approach will offer a route to aid in the interpretation of techniques that generate small interactomes (e.g., affinity purification mass spectrometry and proximity labeling), perhaps even better separating true positives, false positives, true negatives, and false negatives.

Second, we are actively investigating the application of our approach to better understand IDR conservation and functional annotation. We and others have historically leaned heavily on sequence features (e.g., amino acid composition and patterning) to understand conservation in IDRs where primary structure is poorly conserved (*14*, *19*, *20*, *96–102*). In some cases, we anticipate that the conservation of sequence features reflects the preservation of intrinsic biophysical properties of an IDR (*45*, *65*, *103*). In other cases, we anticipate that the conservation of sequence features reflects the conservation of chemical specificity (*14*, *100*). Indeed, the classification of IDRs into distinct chemically similar clusters revealed many IDRs with very different sequences that share similar global interaction fingerprints (**Fig. 5B**). With the methods described here, we now have tools to predict both biophysical properties and chemical specificity directly from sequence, opening the door to new routes for the systematic assessment of conservation in IDRs through the lens of chemical physics (*45*).

In addition to conservation, we see the ability to define context-dependent domains as a potentially important step towards generalized functional annotation in IDRs. In general, we consider protein “domains” to be defined by a function. In practice, the tight link between structure and function for folded proteins has enabled structure to act as a surrogate for function, such that there many examples of conserved domains where – while function remains elusive – we are comfortable referring to them as domains because it is anticipated they will have a stand-alone molecular function in some context. A given IDR may interact with different partners via different regions, such that the definition of a domain is unavoidably context-dependent. Our approach here enables domains to be defined based on intermolecular interaction profiles, enabling subdomains to be defined either with respect to a specific partner or by using a set of precomputed “master chemistries” associated with IDRs (**Fig. 5F**). Indeed, to develop an unbiased route to identify these chemically-distinct subdomains, we clustered all possible dipeptide sequences into 36 chemically related clusters, where clustering was determined by the matrix of dipeptide:dipeptide heterotypic interaction ɛ values (**Fig. S11**). This analysis unveiled a set of chemically orthogonal dipeptide repeats that can be applied to fingerprint an IDR of interest. Using these dipeptides as fiducial markers in chemical space, we can measure the distance between two IDRs based on their difference from these chemical fingerprints.

Third, the throughput and generalizability of our approach lend themselves to the rational design of IDRs with desired intermolecular interaction properties. For example, identifying variants predicted to enhance or suppress phase separation and/or partitioning into an existing condensate with known components becomes trivial. Moreover, rationally designing IDRs that flank binding motifs to assess the role these flanking IDRs have in tuning affinity and specificity is straightforward, as is designing sets of IDRs that modulate intermolecular repulsion in the context of entropic force generation. In short, we see a wide range of design applications here.

### Caveats and limitations

We feel it essential to be explicit and direct about the many caveats and limitations associated with this approach. The *Supporting Information* provides a more extended discussion of these points, but it serves no one to overstate the efficacy, accuracy, and generalizability of a method nor to obfuscate its limitations.

A central assumption in this approach is that the attraction between two IDRs is mediated solely by complementary chemical interactions (chemical specificity), not via precise “structured” interaction between subregions. As such, where sequence-specific interactions will occur, we anticipate false negatives with respect to IDR-mediated interaction. Moreover, for interactions with folded domains, we do not expect to necessarily identify motif binding sites on folded domain surfaces or short linear motifs in IDRs. If anything, we expect sequence-specific binding regions to contribute minimally to chemically-specific attractive interactions, which could reflect competition between a structured bound state and a fuzzy partially bound state. We do not encode mutual information between distal regions, such that the apparent valency between two IDRs is always maximal, regardless of if, in principle, there should be a smaller number of mutually exclusive modes of interaction. Finally, the approach does not consider the intrinsic competition between intramolecular interactions and intermolecular interactions, although this effect can be accounted for explicitly (e.g., see **Fig. 5**).

The predictions made here rely on parameters obtained from coarse-grained molecular mechanics force fields — in this case, CALVADOS (CALVADOS2) and Mpipi (Mpipi-GG) (*42–45*). The two force fields have been well-vetted but also — unavoidably — have limitations. Notably, as discussed previously, Mpipi-GG appears to underestimate aliphatic hydrophobic interactions, whereas CALVADOS2 may underestimate interactions between valine and other hydrophobic amino acids. Similarly, the treatment of electrostatics via a Coulomb potential combined with a Debye-Hückel screening term will clearly fail to recapitulate experimental dependency on salts in higher salt concentrations, where ion activity and Hoffmeister effects dominate (*104–106*). That said, it is worth noting that CALVADOS2 parameterization has shown good agreement in terms of the salt-dependent saturation concentration (*43*). Similarly, the temperature-dependence of the hydrophobic effect is entirely absent from the functional forms of the force fields used here, and while hydrophobic and charged residues would drive lower-critical solution temperature (LCST)-type attractive interactions at higher temperatures, this kind of behavior is not currently captured (*107–110*). Finally, recent work has shown local charge effects can drastically influence the ionization state of charged residues, a phenomenon known as charge regulation, which is not explicitly captured in our model (*111–117*). That said, all of these limitations could, in principle, be parameterized and addressed by altering the underlying model. As such, while at present, these represent limitations, we suggest our approach could be a useful way to improve coarse-grained force fields (e.g., using temperature-dependent peptide solubility to infer temperature-dependent descriptions of intermolecular interactions). Moreover, because of the underlying architecture of FINCHES, if/when new parameters become available (e.g., improved models, new residues, etc.), they can be immediately introduced and used.

The calculation of heterotypic ɛ values should not be in isolation inferred as predicting likely interactomes between IDRs. In **Fig. 4** we calculate many IDRs that have thousands of partners with favorable heterotypic ɛ values. This is not meant to imply that these proteins interact with thousands of partners; each prediction is done in isolation. In reality, the specificity associated with a protein’s interaction profile is a combination of the affinities for partners and the availability (i.e., concentration) of those partners(*15*). If partners are bound by other biomolecules, then their effective concentration is low, even if their absolute concentration is high. As such, individual heterotypic predictions should be treated akin to an *in vitro* experiment - just because two proteins interact *in vitro* does not necessarily mean they interact together *in vivo*, especially if such an interaction is one of many equally favorable interactions observed to seemingly unrelated partners. Nevertheless, if you know two proteins do interact *in vivo*, using *in vitro* experiments to identify regions or residues that may underlie that interaction can be extremely fruitful. This is where we see our approach being particularly useful.

In summary, we want to re-emphasize that our approach here should be seen as an effective route to obtain qualitative (and sometimes semi-quantitative) insight into how an IDR may be interacting with a partner. The impact and consequences of the various caveats considered here should always be considered when assessing whether or not such a prediction is valid or reasonable.

## METHODS

Functional forms of non-bonded terms for Mpipi and CALVADOS force fields were reproduced and implemented in the FINCHES Python package. For Mpipi, the two components here are a Wang-Frenkel potential and a Debye-Hückel potential (*42*, *118*, *119*). For CALVADOS, this is a shifted and truncated Ashbaugh-Hatch potential with a Debye-Hückel potential using an empirical correction for the temperature-dependence of electrostatic interactions (*43*, *120*, *121*). force field parameters for CALVADOS2 and Mpipi-GG were taken from their respective publications (*43*, *45*).

Local charge effects were accounted for by considering the local i+1 and i-1 charge around a charged residue and down-weighting like-charged regions based on local charge density, effectively reducing the repulsion associated with clusters of like-charged residues. Local hydrophobic effects were accounted for my considering contiguous runs of two or three or more aliphatic residues, scaling up aliphatic-aliphatic attractive interactions by 1.5x and 3x for residues embedded within runs of two or three or more aliphatic residues, respectively.

Phase diagrams were calculated using the analytical solution to the Flory-Huggins theory developed and implemented originally by Qian *et al.* (*52*). Solvent-accessible surface areas are calculated using MDTraj (*122*). IDR global dimensions were predicted using ALBATROSS (*45*). Disorder prediction is calculated using metapredict V2-FF (*45*, *123*). Rational sequence designs used for examining homopolymer vs. IDR properties were generated using GOOSE (*124*). Proteome-wide analysis was performed using SHEPHARD, with data obtained from UniProt (*67*, *125*). We make extensive use of previously published experimental data and are indebted to the authors for their previously published careful biophysical and biochemical studies. All sequences reported in this manuscript are defined in **Supplementary Table S7**.

### Implementation and distribution

The ability to predict ɛ-based intermolecular interactions is implemented in our software package FINCHES (**<u>F</u>**irst-principle **<u>IN</u>**teractions via **<u>CHE</u>**mical **<u>S</u>**pecificity). FINCHES not only continues our march toward obscurity with respect to acronyms, but implements both Mpipi-GG and CALVADOS2 modules via a set of common interfaces that enable identical analysis and predictions. Moreover, the underlying architecture makes it straightforward to implement additional force fields in an object-oriented manner. FINCHES is fully open source and available at https://github.com/idptools/finches.

To facilitate adoption and use, we also provide several Colab notebooks that enable standard types of analysis (intermaps, phase diagram prediction, etc). These are linked from https://github.com/idptools/finches-colab.

## ACKNOWLEDGEMENTS

We thank Shahar Sukenik, Broder Schmidt, Max Staller, and members of the Holehouse lab for feedback and discussion. We thank Kresten Lindorff-Larsen and Sören von Bülow for being willing to - once again - coordinate preprint submission. We are indebted to the various groups who provided high-quality data with which we could directly test predictions. We also extend a warm thank you to Jerelle Joseph, Aleks Reinhardt, Rosana Collepardo-Guevara, Giulio Tesei, and Kresten Lindorff-Larsen for their prior and ongoing work on the Mpipi and CALVADOS forcefields.

This work was funded by the NIH (DP2-CA290639 to A.S.H.), an NSF GRFP fellowship (DGE-2139839) to J.M.L., a Keck Postdoctoral Fellowship to E.T.U., and a MilliporeSigma Fellowship to G.M.G.

