## Supporting Information for "Direct prediction of intermolecular interactions driven by disordered regions"

**VERSION 1.0 (June 3rd 2024)**

### Extended methods

#### Implementation

FINCHES is fully open source and hosted at https://github.com/idptools/finches. It is implemented in Python (https://www.python.org/) (version 3.7 or higher) with Cython implementations for a subset of performance-sensitive algorithms (https://cython.org/). FINCHES also makes extensive use of NumPy, SciPy, and Matplotlib (*1–3*). Version control is provided via git (https://git-scm.com/). Bugs and features can be reported/requested at https://github.com/idptools/finches/. Additional forcefields can be implemented in the `finches.forcefields` module.

The finches-online webserver (http://finches-online.com/) is built using Flask (https://flask.palletsprojects.com/) and uses nginx (https://nginx.org/en/) and gunicorn (https://gunicorn.org/) for backend infrastructure. It provides front-facing services for intermap construction and phase diagram prediction.

#### Analysis

Analysis for this manuscript rely on metapredict, SHEPHARD, ALBATROS and sparrow (*4–7*). The figures and analysis associated with this manuscript are fully reproducible via Jupyter notebooks shared at https://github.com/holehouse-lab/supportingdata/tree/master/2024/ginell_et_al_2024.

#### Calibration of charge prefactor

We implemented a simple charge correction that downweighs repulsion from clusters of like-charged residues to capture local charge effects. We sought to implement the simplest charge correction possible to capture local charge effects. This goal was motivated by the fact that more complex corrections would require more parameters and decisions, and – at this stage – our goal is to implement an approach that uses the native forcefields as closely as possible.

The local charge correction is implemented when calculating the initial intermolecular ϵ matrix (i.e., without any sliding window analysis). If two residues in opposing chains are both charged, we excise the i+1 and i-i residues around the two residues in question and concatenate these two subsequences together to generate a fragment with a maximum length of six amino acids (minimum of four, in the case of two end residues). For that fragment, we calculate the fraction of charged residues (FCR), which lies between 0 and 1, and the net charge per residue (NCPR), which lies between -1 and +1. The weighting factor is then computed as |NCPR/FCR|. This means that for fragments of all one type of charge, the weighting factor is 1 (1/1), whereas for net neutral fragments, the weighting factor is 0 (0/1). Intermediate values are then tuned based on the fragment's FCR and NCRP values. The weighting factor is applied to each instance where pairs of residues in opposing chains are both charged. Finally, all those weighting factors are scaled in a forcefield-specific manner via a single scalar we refer to as the *charge prefactor*. The charge prefactor is the only free parameter in our charge correction, and allows different force fields to implement differing levels of local charge effects. The charge prefactor must lie between 0 (no local charge effects) and 1 (weighting factors used directly).

We calibrated the Mpipi-GG charge prefactor at 0.2 based on tuning the self-interaction propensity of the Das-Pappu sequences (**Fig. S16**) (*8*). Briefly, based on single-chain compaction results alongside work from Lin and Chan, we anticipate sequences with large blocks of oppositely charged residues (i.e., high-kappa sequences) to have a negative (attractive) ϵ in the zero-salt limit (*9*). A charge prefactor of 0.2 defines the boundary between attractive and repulsive ϵ values for these sequences in a reasonable location, given previous theoretical predictions and simulation work (**Fig. S17A**)(*9–11*). For CALVADOS2, the charge prefactor was calibrated to match the (magnitude-corrected) trends described by Mpipi-GG, giving a charge prefactor of 0.7 (**Fig. S17B, C**). This calibration is relatively empirical and qualitative. While a more robust calibration is certainly possibly reached, we suggest attempting to fit a more precise dependency here may over-extend the reasonable predictive power of the underlying force field, especially given the layers of simplifying assumptions being made.

**Aliphatic residue clustering**

Aliphatic residues also employ a local correction factor. Specifically, for contiguous runs of two, three, or more aliphatic residues, the aliphatic-aliphatic attractive interactions are scaled by 1.5x and 3x for residues embedded within runs of two or three or more aliphatic residues, respectively. Given the limitation in coarse-grained models, we developed his empirical correction motivated by the realization that aliphatic residues can form dry, stable interfaces if a sufficient number of aliphatic residues are together. However, this effect is a consequence of the discrete nature of water molecules and – we reasoned – should be explicitly introduced for a coarse-grained implicit solvent mode.

**Comparison of Mpipi-GG and CALVADOS2-based predictions**

We note that almost all results in this manuscript are reproduced by both CALVADOS and Mpipi-GG-based predictions. The singular exception to this is the impact of the conserved region (CR) in TDP-43 (**Fig. S6**). For these, Mpipi-GG-based predictions do not correctly capture the hydrophobic nature of the aliphatic residues in the CR, whereas CALVADOS2 does a good job. To reiterate, neither model has any structural knowledge, so identifying this hotspot reflects the local density of aliphatic residues.

Mpipi-GG and CALVADOS2-based predictions are generally reported in their natural units, which place CALVADOS2 scores ~4x larger than Mpipi-GG scores. The exception to this is in **Fig. 5C** and **Fig. 5F**, where Mpipi scores are multiplied by a scalar to better overlap with the CALVADOS2 scores - this is solely for visual purposes, in that numerical comparison of absolute values between the two models is not relevant or informative.

**Intermap construction**

Intermaps are calculated using a sliding window between all possible intermolecular fragments of a specified size. In general, for larger IDRs, we default to a size here of 31 residues, although for smaller IDRs, this leads to a loss of fine-grain detail. For example, intermaps associated with the analysis in **Fig. 5** use a 13-residue sliding window.

**Clustering of IDRs in chemical space (human proteome)**

Chemical clustering of natural IDRs (**Fig. 4**) was performed by calculating all possible heterotypic $\epsilon$ values between all 3,413 IDRs in the human proteome that are between 100 and 150 residues in length. This consists of 11,648,569 individual $\epsilon$ calculations, generating the complete non-redundant matrix of intermolecular $\epsilon$ scores. Having generated this matrix, we use Ward's method for hierarchical clustering, followed by the optimal ordering of the resulting clusters to minimize the distance between adjacent clusters in chemical space. We used 24 clusters based on varying the number of clusters to assess the extent of chemical decomposition. We arrived at 24 clusters as a reasonable number that provided chemically interpretable and distinct groups. Cluster assignment for each IDR is provided in **Supplementary Table S4**.

**Clustering of IDRs in chemical space (chemical fingerprints)**

Chemical clustering of all possible dipeptides (**Fig. S11**) was performed by calculating all possible heterotypic $\epsilon$ values between all 210 non-redundant dipeptides. We used a repeat of 20 amino acids (10 repeats) for $\epsilon$ calculations. As before, having generated the matrix of all intermolecular $\epsilon$ values, we used Ward's method for hierarchical clustering, followed by the optimal ordering of the resulting clusters to minimize the distance between adjacent clusters in chemical space. We specified 36 clusters for dipeptide fingerprint chemical decomposition based on varying the number of clusters and observing when intuitively consistent chemical groups emerged. Full cluster memberships are provided in **fingerprint_clusters_calvados.fasta** and **fingerprint_clusters_mpipi.fasta**, while one member-per-cluster with representative members (as shown in **Fig. S11**) are provided in **fingerprint_mpipi.fasta** and **fingerprint_calvados.fasta**.

**Phosphomimetic mutations**

Experimentally verified phosphosites were taken from ProteomeScout and parsed using SHEPHARD, as described previously (*5*, *12*). For every site, we ensured the corresponding position matched the expected unmodified residue (i.e., pSer = Ser, pThr = Thr, and pTry = Tyr) and then converted every phosphosite to glutamic acid (E). In total, this involved 130,606 phosphoserine sites, 53,744 phosphothreonine sites, and 38,106 phosphotyrosine sites. We note that a glutamic acid phosphomimetic likely does not fully capture the physical chemistry associated with phosphorylation, but previous work examining phosphomimetic mutations in the context of phase separation suggests these are reasonable approximations (*13*, *14*). This analysis was performed over all IDRs without length filtering.

**Protein copy number analysis**

Protein copy number information was taken from quantitative mass spectrometry analysis on HeLa cells performed by Hein et al. (*15*).

**Gene Ontology (GO) analysis**

Gene ontology (GO) analysis was performed using PANTHER-DB (release 2024-02-26, 10.5281/zenodo.10536401 Released 2024-01-17) (*16*). Enrichment analysis was performed in terms of biological processes with P-values computed based on Fisher's exact test, and multiple hypothesis correction was done in terms of false discovery rate (default settings). This analysis was performed over proteins where one (or more) IDR was 100 residues or longer.

For Mpipi-GG and CALVADOS2 enrichment analysis, we compared proteins with an attractive $\epsilon$ value ($\epsilon$ <0) with all proteins that have an IDR over 100 residues. This is the appropriate background to compare against to answer the question, *"For proteins with long IDRs, what types of biological processes are enriched for when considering those IDRs that are homotypically attractive*?".

For WordCloud generation, we filtered GO hits with the following criterion: over 40 different proteins must have been associated with the label, p-value of less than 0.001, and enrichment of over 1.25 vs. the background. The WordCloud was generated using the wordcloud Python package (https://pypi.org/project/wordcloud/).

### Extended discussion

**How to interpret phase diagrams**

A phase diagram can be thought of as a map. Just as geographical maps describe the local landscape at some latitude and longitude, a phase diagram describes the state of matter at some latitude (x-value) and longitude (y-value), where the x and y axes define the phase space being examined. In our example in **Fig. S4A**, that phase space is temperature and protein volume fraction (i.e., protein concentration).

The phase space associated with the phase diagram shown in **Fig. S4A** is divided into two distinct regimes - the two-phase regime and the one-phase regime. If the system finds itself at a temperature and concentration in the one-phase regime, thermodynamics will favor the protein being uniformly distributed. This says nothing about the concentration of the protein - i.e., the protein could be uniformly distributed and very high in concentration. If the system finds itself at a temperature and concentration in the two-phase regime, thermodynamics will favor the protein demixing into two coexisting phases: a dense phase and a dilute phase.

The top of the phase diagram is defined by a critical point, set by the **critical temperature** ($T_c$) and the critical concentration or **critical volume fraction** ($\Phi_c$). $T_c$ is the temperature above which the system will never be in the two-phase regime, regardless of the protein concentration. $\Phi_c$ defines the concentration where dense and dilute phases converge. The **binodals** define the boundary between the two-phase and one-phase regime, while the **spinodals** define an inner boundary that delineates the kinetics of phase separation.

Measuring full-phase diagrams is generally laborious and challenging - for more details on this, see reviews by Posey et al. and Peran et al. (*17*, *18*). Instead, it is often more common to measure the **saturation concentration** ($C_{sat}$) as a function of temperature, which reports on the part of the low-concentration arm of the binodal. $C_{sat}$ can be directly measured as the concentration of protein remaining in the dilute phase (supernatant) after phase separation has occurred or can be inferred using a binary droplet assay to define a concentration boundary across which droplets are first observed. In general (for two-component systems that show upper critical solution temperature [UCST] behavior), the $T_c$ and saturation concentration are well-correlated and report on the strength of the cohesive interactions that drive self-association.

**Inter-residue ϵ *vs. inter-residue $B_2$* (second virial coefficient)**

ϵ is a mean-field parameter quantifying the attractive or repulsive interactions between two components. In the limit of a system comprising exactly two isotropic spherical components (e.g., two spherical beads with an isotropic interaction potential in a box), ϵ values should be directly proportional to the second virial coefficient. Indeed, considering the Mayer-f function involves taking an integral over the inter-residue interaction potential with a volume element correction (**equation 1**), it should be unsurprising that calculating an analytical $B_2$ (possible in the limit of the only degrees of freedom being inter-bead distance, which is the situation for a two-bead system with an isotropic potential) reveals a near 1:1 agreement with ϵ (**Fig. S18A**).

$$B_2 = 2\pi \int 1 - e^{-u(r)/kT} r^2 dr \quad (1)$$

The small differences between $B_2$ and ϵ reflect the fact that the ϵ calculation does not factor in a volume-element correction (discussed below). If we recompute the second virial coefficient and do not consider the volume element (i.e., remove $r^2$ from equation 1), the resulting ϵ and $B_2$ values are directly proportional (**Fig. S18B**). This is neither profound nor surprising, but we include it here for completeness.

At the level of inter-residue ϵ values, the difference between $B_2$ and ϵ is that $B_2$ considers the volume element around the bead, whereas ϵ makes no such 3D volume correction. To make this more concrete, let us imagine a scenario where we have two pairs of beads with identical shapes and areas associated with their inter-residue interaction potentials, yet for one of those pairs, one of the beads is much larger than the others. Suppose a volume correction was used when calculating $B_2$ (as it is normally). In that case, the pair with the larger bead will have a larger $B_2$, yet if the areas associated with the potentials are the same in both cases, they would have identical ϵ values (**Fig. S18C**). We deliberately chose to discard the volume correction here and use the naive integral area because a core component of our underlying assumption is a true mean-field approximation in that no excluded volume or chain connectivity is considered. To include a bead volume component in the calculation would be inconsistent with that approximation, such that we effectively remove bead size as a factor from our calculation.

**The decision to report ϵ based 'scores' as opposed to a unit-based quantitative metric**

As discussed above, one could factor in a volume element and convert the underlying ϵ values into units of kcal/mol. However, we have deliberately decided to report ϵ values without units and as qualitative "scores" instead of quantitative values. This decision is partly motivated by the desire to avoid implying a degree of accuracy, precision, or comparison to real-world measurable phenomena. As discussed below (*On the utility of* ϵ*-based predictions*), the value of an approach like this is not in obtaining quantitative predictive values but in quickly and easily acquiring intuition as to likely determinants of molecular function, where that intuition is derived from a quantitative and interpretable source.

**On the utility of ϵ-based predictions**

We have done our best to balance highlighting the value in an ϵ-based analysis and laying out the caveats and limitations. We fully accept that there are likely additional caveats and limitations we have failed to explicitly enumerate (hopefully, we can include those during peer review). Nevertheless, we anticipate this type of analysis (repurposing force field chemical physics for informatic analysis via a common framework, such as FINCHES) will open the door to novel integrative high-throughput bioinformatic analysis.

As is always the case, the resolution of a model defines the resolution of the questions that a model can reasonably answer. Here, we have deployed a low-resolution model (in structural terms). With this in mind, we see predictions that emerge from ϵ-based analysis as offering two types of insight. When analyzing large datasets, these predictions provide a route to uncover general trends driven by chemical specificity. When focusing on a specific protein or a small number of proteins, these predictions provide a route to motivate hypothesis-driven experiments or interpret experimental results through the lens of chemical specificity. In both cases, chemical interpretability sits at the heart of these predictions.

Compared to machine-learning-based approaches, we propose that physics-based bottom-up models offer certain advantages for developing specific testable hypotheses and understanding the molecular origins of trends identified from large datasets.

First, the quality of a machine learning model depends entirely on the quality of the available training data. At present, the lack of experimentally derived high-quality intermolecular interaction data obviates the value of *bona fide* machine learning models for IDR-associated interactions. That said, even with small datasets, simple models may offer valuable qualitative insight (*8*, *19–21*). An alternative strategy is to use molecular simulations to generate large volumes of high-quality data for surrogate model training, a strategy we (*6*) and others (*22*, *23*) have taken. While such an approach benefits from the guarantee of homogeneous and essentially unlimited training data, a current limitation is that the underlying predictions are constrained to the conditions under which simulations were performed. This limitation is not unlike any other machine learning scenario, but given that IDR-mediated interactions are often environmentally sensitive, the combinatorics of building appropriate training sets across a range of environmental conditions may be prohibitively challenging. In contrast, given our predictions originate from an interpretable physical model, that underlying model can – in principle – be tuned rationally. In this work, we highlight this by altering the ionic strength and illustrating how this tunes intermolecular interaction strength (both weakening [**Fig. 3F, Fig. S12**] and strengthening [**Fig. S13]** in a sequence-dependent manner). However, in principle, many additional environmentally-dependent modulations could be encoded in the underlying force field physics, such as temperature-dependent hydrophobicity, solvent-dependencies, etc.

Second, model predictions may be wrong. How often a prediction is wrong depends on the model's predictive power. A shortcoming of machine-learning-based predictions is that when predictions are wrong, we generally don't know *why* they are wrong. This simple reality makes it difficult for humans to learn from a model's inaccuracies and extrapolate them to other predictions. It also makes it difficult to identify places where our underlying assumptions may be wrong.

In contrast, our approach's interpretability enables us to understand the underlying chemical logic for each prediction. This makes it possible to understand failure modes and recalibrate our intuition and application of such a model. It also provides a route to understanding limitations and motivating further improvements in specific aspects of force field chemistry.

Finally, we focus here on using coarse-grained force fields. However, one could – in principle – empirically calculate inter-residue mean-field potentials for higher-resolution models (i.e., umbrella sampling every pair of residues to calculate an apparent inter-residue potential energy profile). It is unclear whether such an approach would offer any advantages over lower-resolution models because almost all the advantages a high-resolution model offers would be discarded when constructing an empirical potential. However, for elucidating direct environmentally-responsive behaviors for complex solvents, high-resolution models that enable the effective mean-field consequence of those environments on the inter-residue potential of mean force calculations could offer a route to 'coarse grain' in chemical space.

**ϵ-based predictions vs. explicit simulations**

One important point we wish to emphasize is that our approach here in no way relegates explicit molecular simulations to second-class citizens. In fact, for both CALVADOS and Mpipi, we see our approach as complementary in that it offers a rapid way to prototype expected intermolecular interaction profiles when considering solution conditions or sequence changes. Furthermore, the simplifying assumptions made here (e.g., discarding spatial information and correlated interactions, see below) are precisely the types of insight simulations provide). We would also welcome the integration of additional molecular force fields into FINCHES.

**Limitations of mean-field approximations for intermolecular interaction**

Some analysis presented here uses an overall single mean-field value (ϵ) to predict phase diagrams. Mean-field descriptions simplify the complex interactions of a many-body system by approximating the effect of all components on any given particle as an average (mean) effect. Mean-field models have found great utility in the context of phase separation, implicit solvent models, and in capturing solute effects on biomolecules (*24–26*). However, mean-field descriptions also introduce limitations - notably, they fail to capture correlations between components over which the field is calculated and do not appropriately deal with fluctuations (e.g., around the critical point of a phase diagram). The fact a mean-field-based approach does a reasonable job of predicting simple homotypic phase behavior is consistent with the fact that single-chain dimensions are often (although not always) well-correlated with phase behavior for homotypic systems (*27*, *28*). In both scenarios, a single value captures the average intramolecular interactions, an underlying component of Flory-Huggins theory.

While a mean-field value may be reasonable for capturing phase behavior in some scenarios, we do not expect ϵ to correlate 1:1 with the second virial coefficient ($B_2$). This expectation is illustrated in recent work by Chen & Jacobs, where a modest correlation of  0.67 was found between directly computed second virial coefficients (from simulations) vs. the inferred virial coefficient obtained based on Flory-Huggins theory (*29*). In contrast, the mean-field approach works remarkably well for Chen & Jacobs in predicting phase diagrams. With this in mind, we suggest major caveats should be considered if using ϵ to optimize for intermolecular interactions.

In short, ϵ-based predictions should generally be interpreted as qualitative descriptions of the types of interactions that could happen instead of hard predictions of things that will happen. In our experience, when a subregion is predicted to interact strongly with another subregion, that is likely a genuine interaction. If a subregion is not predicted to interact, that could be a false negative since two

IDRs may fold together (or one IDR folds to bind the surface of a folded domain), modes of interaction not captured in our approach. Rather than a weakness, we suggest this offers a route to diagnose likey sequence-specific interactions - if an IDR empirically binds a partner yet there is no obvious complementary chemistry between the two in either Mpipi-GG or CALVADOS2-based ϵ predictions, this would imply a structural interaction is taking place.

***Physics-based models in a deep-learning world***

Over the last five years, deep learning has revolutionized protein biophysics, with AlphaFold2, RoseTTAFold, and AlphaFold3 (and derivatives thereof) setting the bars for protein structure and complex prediction (*30–33*). Deep learning-based approaches necessitate large volumes of high-quality training data to learn the associated mapping between input and output, ideally where a 1-to-1 mapping between input and output exists. Given these constraints, protein structure prediction is, in some ways, a perfect problem for deep learning. The Protein Databank offers an exceptionally well-curated, high-quality dataset with (generally) a defined and deterministic sequence-to-structure mapping that is largely insensitive to context. In contrast, IDR-mediated intermolecular interactions lack many features that facilitate the efficacy of deep learning. IDRs lack an extensive repository of high-resolution training data; any such data would be inherently context-dependent, and no single experiment offers a 'silver bullet' to map from sequence to interaction.

There has been recent interest in using machine learning to predict phase separation from sequence. These tools are typically trained on sequence datasets with <1000 sequences where phase separation/condensate formation has been observed *in vitro* and/or in cells, yielding methods that predict the propensity of a protein or protein region to undergo phase separation. Beyond the mismatch of the number of data points vs. model parameters, the lack of a true negative dataset, the inherent bias in the types of proteins for which we have phase separation data, and the conflation of *in vitro* phase separation *vs.* in-cell condensate formation/association, we see two conceptual problems with this approach.

First, training on data that allows concentration, temperature, and additional cosolutes to be uncontrolled for makes this a near-uninterpretable learning problem. Absent constraints on these three variables, the answer to "Will my protein phase separate?" is simply, "Yes." Second, asking whether a protein undergoes phase separation is an inherently under-defined question in that it misses specifying with whom. If this is a question of homotypic phase separation, it bears thinking as to whether this is a physiologically relevant question. If this is a question of heterotypic phase separation, then the identity of the partner(s) is essential. As such, we remain skeptical of the utility of machine learning methods in this space without robust training data that, at the very least, titrate the total concentration of a large number of sequences to obtain saturation concentrations under fixed solution conditions.

Von Bulow et al. recently directly addressed the challenge of machine-learning-based prediction of homotypic phase separation (*22*). This work uses a combination of active learning and molecular dynamics simulations followed by a machine learning protocol to train a homotypic phase separation predictor. The resulting method circumvents many of the issues described above. It enables highly accurate predictions of saturation concentrations concerning what a CALVADOS2 model would predict, which shows good predictive power compared to extant *in vitro* experiments. While such an approach is limited to homotypic interactions under the conditions parameterized, it circumvents almost all of the issues raised for experimentally-based training of machine learning models, and we are bullish that this will be a valuable contribution to the field.

### Supplementary tables

**Supplementary Table S1:**

Homotypic ϵ predictions using Mpipi-GG for human IDRs above 100 amino acids in length (9,909 proteins)

**Supplementary Table S2:**

Homotypic ϵ predictions using CALVADOS2 for human IDRs above 100 amino acids in length (9,909 proteins)

**Supplementary Table S3:**

19,702 human IDRs showing the change in homotypic ϵ upon complete phosphorylation. A and B for CALVADOS2 and Mpipi-GG, respectively.

**Supplementary Table S4:**

Assignment of all human IDRs between 100 to 150 residues (n=3,414) to one of the 24 chemical clusters shown in Fig. 4B.

**Supplementary Table S5:**

The average chemical properties for sequences in each of the 24 chemical clusters, as shown in Fig. 4B. Clusters from CALVADOS2 and Mpipi-GG, respectively (note we do not expect 1-1 mapping between clusters).

**Supplementary Table S6:**

Assessment of all human IDRs between 100 to 150 residues that have at least one attractive interactor (n=3,413) ranked by number of attractive interactors. Attractive interactors are proteins for which, in a binary interaction, we would expect favorable mean-field interaction, not that we expect each IDR in this list to interact with the number of listed interactors simultaneously.

**Supplementary Table S7:**

All sequences used in this manuscript

### Supplementary figures

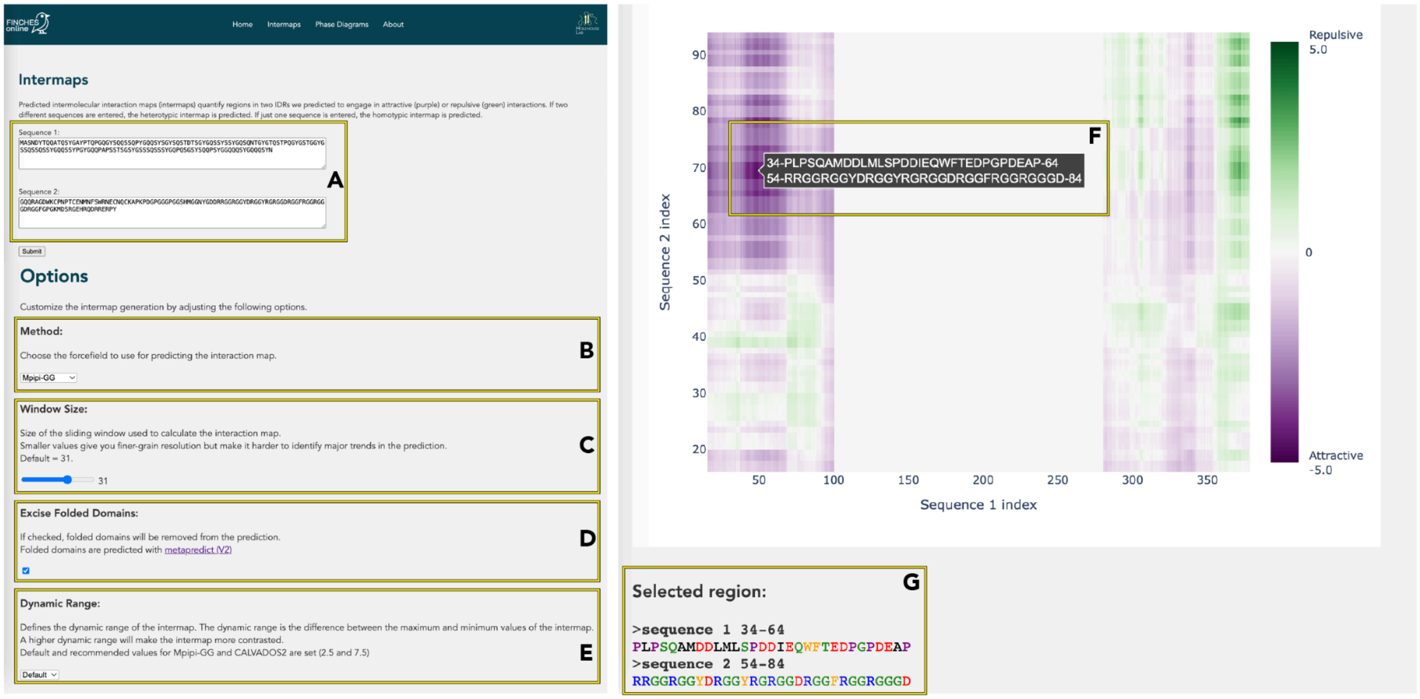


**Fig. S1 FINCHES webserver.** The FINCHES-online webserver (www.finches-online.com) provides easy access to predict intermaps and phase diagrams. The left-hand side of the figure above reports on the input information, which includes **(A)** one (homotypic prediction) or two (heterotypic prediction), **(B)** Forcefield method to be used (Mpipi-GG or CALVADOS2), **(C)** Windowsize over which analysis should be performed (default is 31), **(D)** The ability to toggle on/off folded domain excision, and **(E)** defining the dynamic range over which the prediction shows. Upon intermap prediction, an interactive intermap is generated (right). Hoving the most over a specific region will display the intersecting sequence regions used to obtain the value at that location **(F)**, and clicking on that point will display the two sequence regions colored **(G)** and copy those two sequences to the users' clipboard. In this way, it becomes simple to extract specific regions from the intermap to provide.

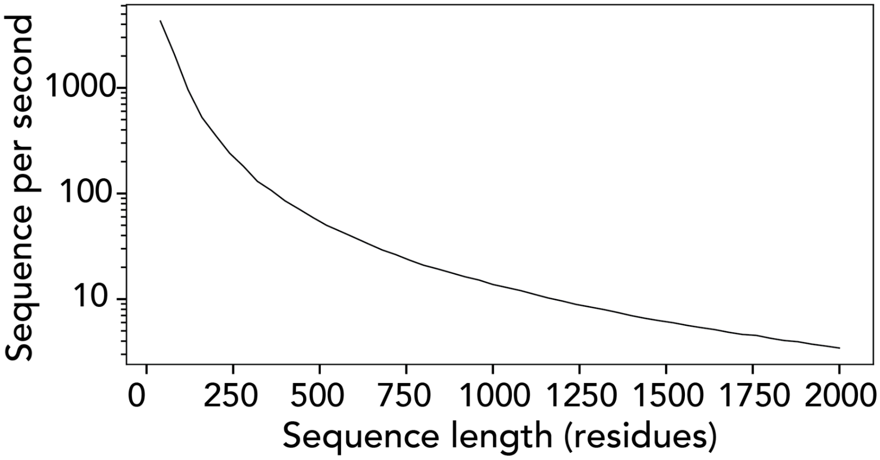


**Fig. S2 Performance for calculating inter-residue ϵ values**

Benchmark performance information using FINCHES to calculate inter-residue ϵ values. For a 100-residue IDR, performance is more than 1000 predictions per second on commodity hardware (Macbook Pro).

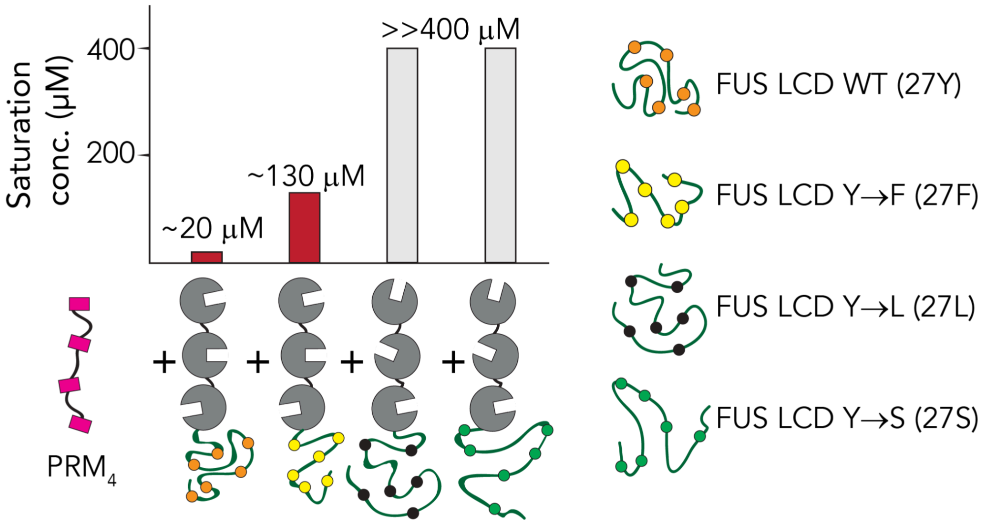


**Fig. S3 Overview of the experimental system used for $A_2$ measurements**

A two-component tandem repeat of proline-rich motifs (PRMs) with a tandem SH3 scaffold is used to measure $A_2$ values. PRM and SH3 bind one another tightly, and $PRM_4$ + $SH3_3$ will phase-separate together (*34*). By using this scaffold as a baseline, Lin & Currie assessed how adding an IDR to the tandem SH3 construct altered the light scattering second virial coefficient ($A_2$). The saturation concentration for $SH3_3$ + $PRM_4$ is around 160 μM, so adding FUS (WT) and FUS (YtoF) reduces that saturation concentration, suggesting these add net attractive interactions. In contrast, FUS (27L) and FUS (27S) impede phase separation, highlighting that the FUS sequence, absent aromatic residues, acts as a solubility tag.

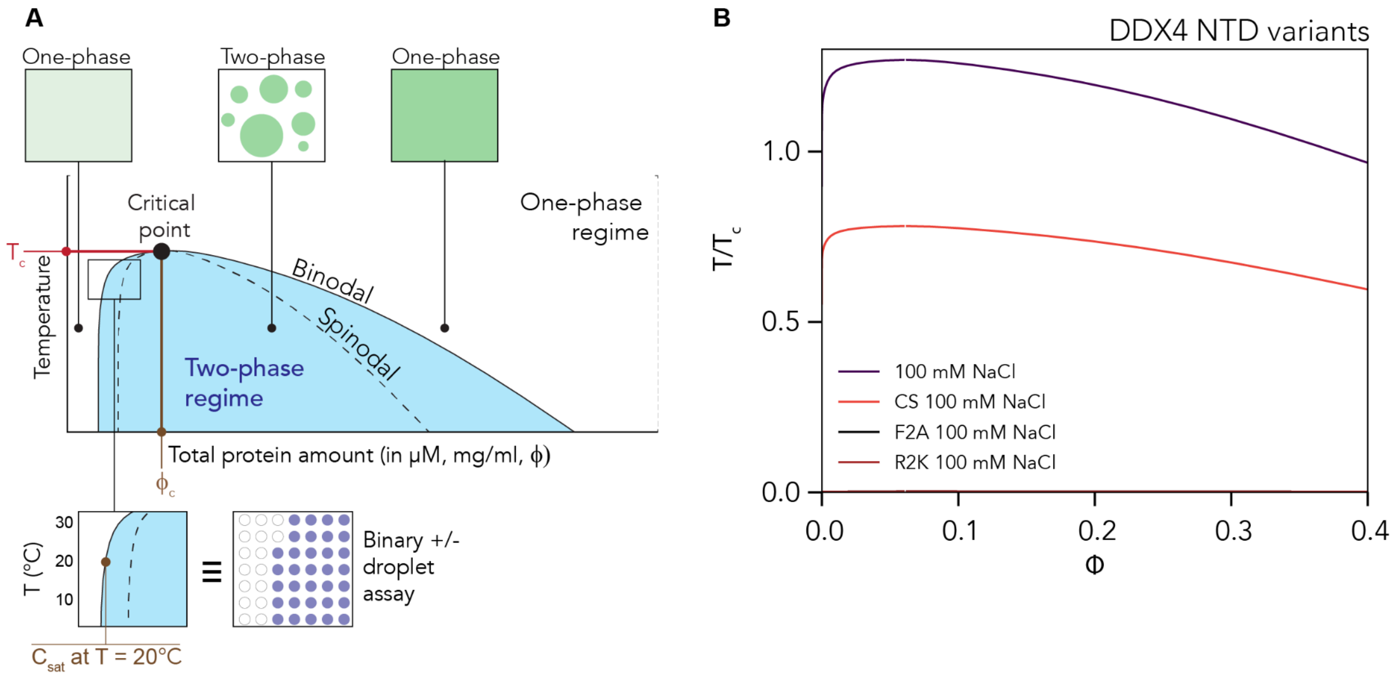


**Fig. S4 Phase diagrams quantify intermolecular interaction (A)** Shown is a schematic and annotated example of a hypothetical phase diagram for a simple two-component system (protein + solvent). A detailed description of how to read and interpret a phase diagram is included in the supplementary methods. **(B) Predicted phase diagrams for four variants of DDX4-NTD at 100 mM salt.** While WT and CS show two-phase behavior, F2A and R2K variants are predicted to be entirely soluble (i.e., no curve is observed), in good agreement with experimental characterization(*35*, *36*).

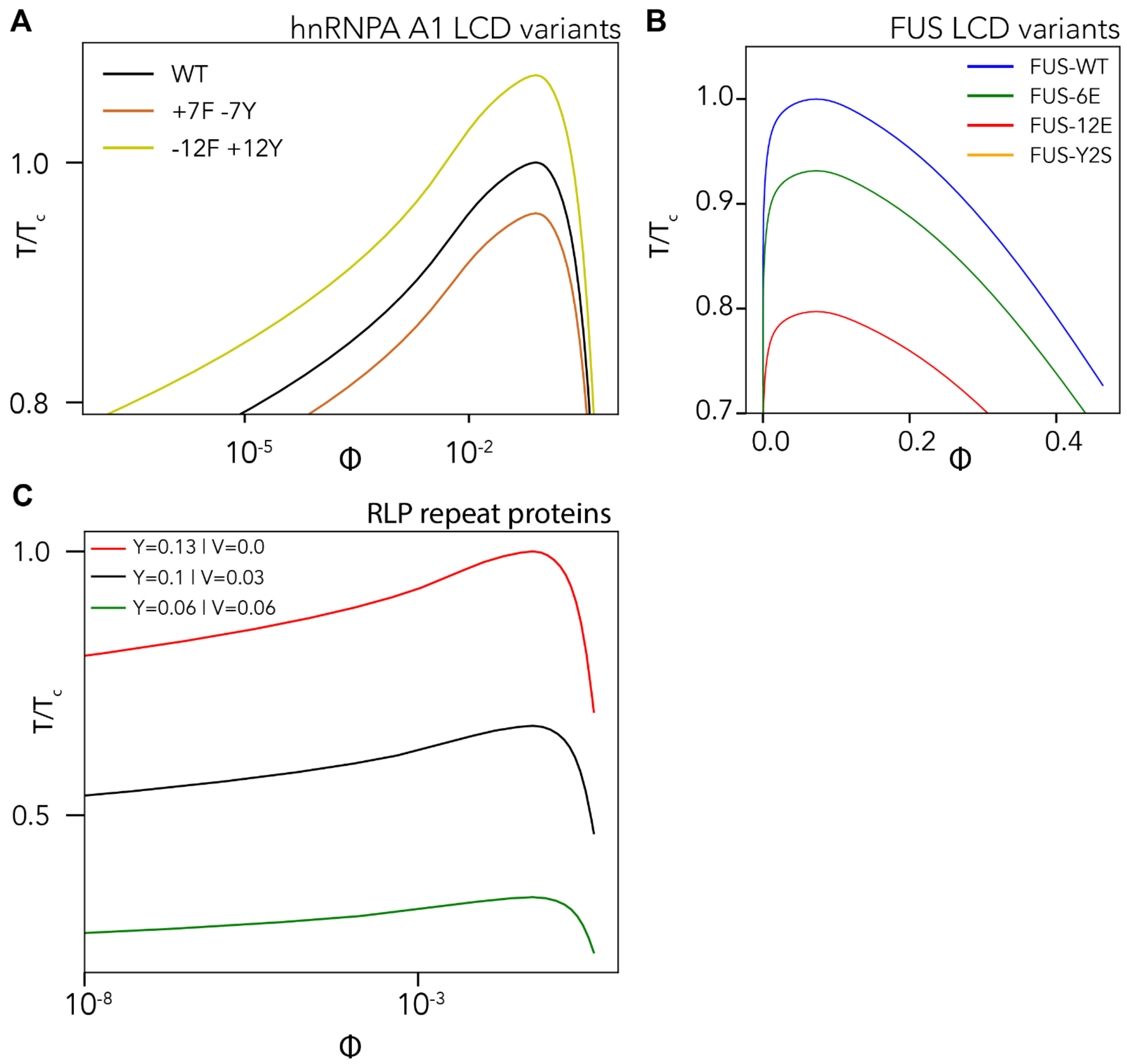


**Fig. S5: Predicted phase diagrams for various experimentally-measured disordered proteins.**

**(A)** Varying the aromatic residues identity in the hnRNPA1 low complexity domain recapitulates trends and relative critical temperatures measured experimentally by Bremer & Farag & Borcherds et al. (*37*). The order of aromatic residue strengths is consistent here in other work, ranking tryptophan (W) > tyrosine (Y) > phenylalanine (*38*, *39*). **(B)** Phase diagrams for the FUS low-complexity domain alongside variants with different numbers of phosphomimetic mutations. These results recapitulate the previously measured impact of phosphorylation and phosphomimetic mutations on FUS (*14*, *40*). Note that FUS-Y2S cannot phase separate at all, consistent with experimental measurements reported by Lin & Currie et al.(*41*). **(C)** Dependence of RLP proteins as aromatic residues are exchanged for valine, recapitulating results reported by Dzuricky et al. (*38*).

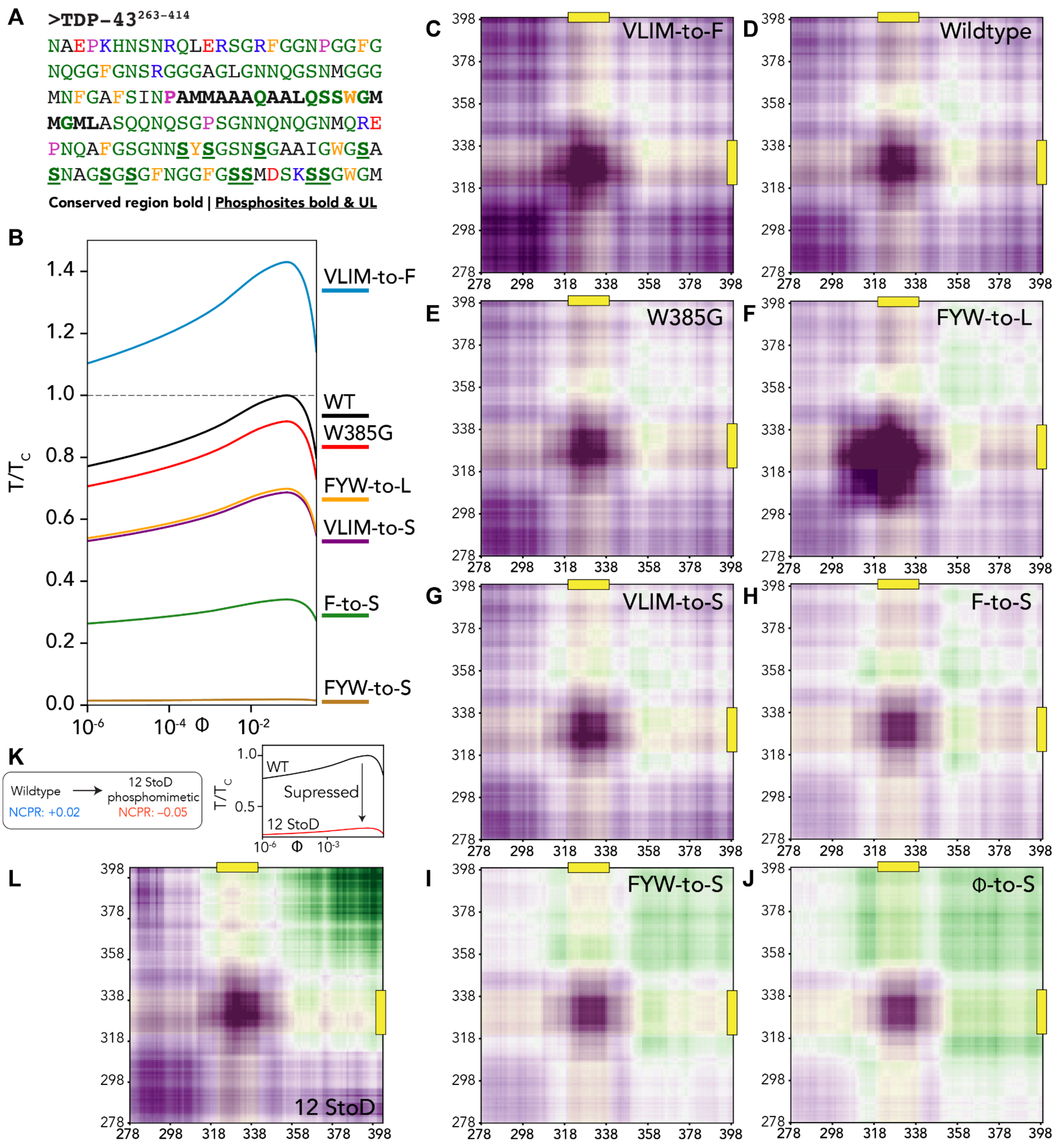


**Fig. S6: Predicted phase diagrams and intermaps for TDP-43 low complexity variants. (A)** TDP-43 low complexity domain (LCD), defined here as residues 263-414, as done by Schmidt et al. (*42*). The TDP-43 LCD contains a conserved region (CR) which forms a hydrophobic alpha helix (bolded, PAM….GML]) along with several C-terminal phosphosites that have been investigated by Gruijs da Silva et al. (*13*). **(B)** Using FINCHES with CALVADOS2, we predicted WT-normalized phase diagrams for a series of variants, recovering experimentally measured trends reported previously (*42*). (**C - J**) Intermaps for all variants reported illustrating how different mutations rewire the predicted intermolecular interactions. The TDP-43 LCD self-interactions have been proposed to be driven by various modes of interaction. Our predictions recapitulate this general phenomenon, illustrating how changes to different chemistries or regions can influence predicted phase behavior and intermolecular interactions. **(K)** Predicted phase behavior for the 12 StoD variant of the TDP-43 LCD recapitulates behavior reported through *in silico*, *in vitro*, *in cell* work by Gruijs da Silva et al. (*13*). **(L)** Predicted intermap suggests that 12 StoD variant drastically weakened the C-terminal half of TDP-43. In contrast, the N-terminal half remains relatively wildtype-like, yet this is sufficient to suppress phase separation dramatically.

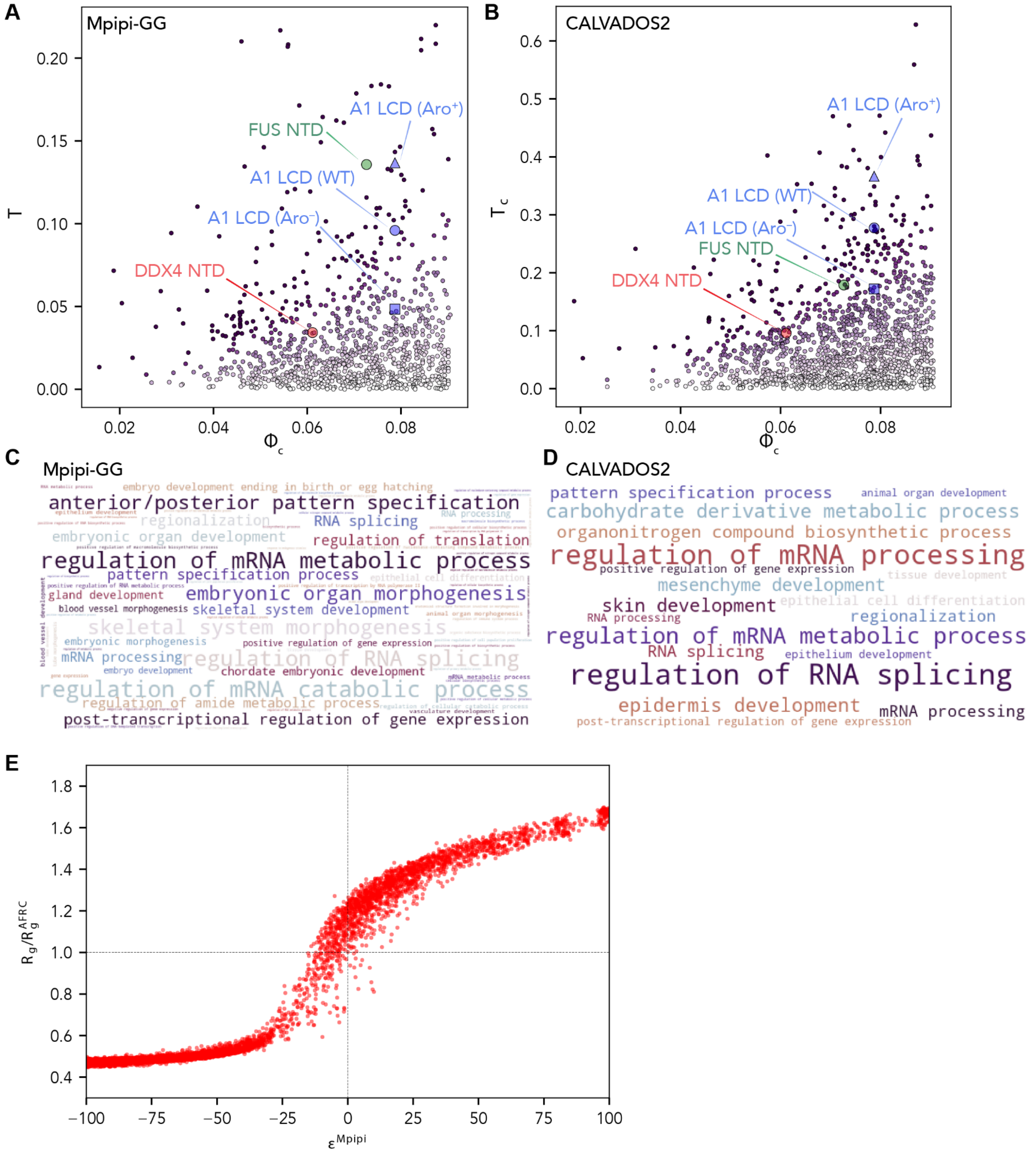


**Fig. S7 Homotypic ϵ analysis. (A, B)** Predicted critical temperature ($T_c$) and critical volume fraction ($\Phi_c$) for human IDRs that are over 100 amino acids in length and have an attractive ($\epsilon < 0$) mean-field interaction. For a description of $T_c$ and $\Phi_c$, see **Fig. S4**. Points are colored by ϵ value (more purple = more attractive). Previously characterized proteins that undergo phase separation *in vitro* have been included for comparison (see **Fig. 3**). Proteins with a high $T_c$ and low $\Phi_c$ are predicted to phase separate the most robustly. (**C, D**) Gene Ontology (GO) analysis for proteins in panels A and B compared to all human proteins with one or more IDR that is longer than 100 amino acids. For both Mpipi-GG and CALVADOS2-derived analyses, RNA-associated biological processes are strongly enriched. **(E)** Synthetic homopolymeric IDRs were generated using GOOSE, and the normalized predicted radius of gyration was compared to the $\epsilon$ value. The radius of gyration was predicted using ALBATROS, a deep-learning-based sequence-to-ensemble prediction tool trained on Mpipi-GG simulations(*6*). This value was normalized using an Analytical Flory Random Coil (AFRC) derived radius of gyration (*43*). The AFRC provides a sequence-specific null model for the expected dimensions of an IDR if the amino acid chemistry had no impact on the underlying ensemble (i.e., if it behaved as a Gaussian chain). This reveals the expected coil-to-globule transition, where $\epsilon$ effectively captures a mean-filed self-interaction potential, and the radius of gyration ($R_g$) reports on the biophysical consequences of that potential in terms of average ensemble dimensions.

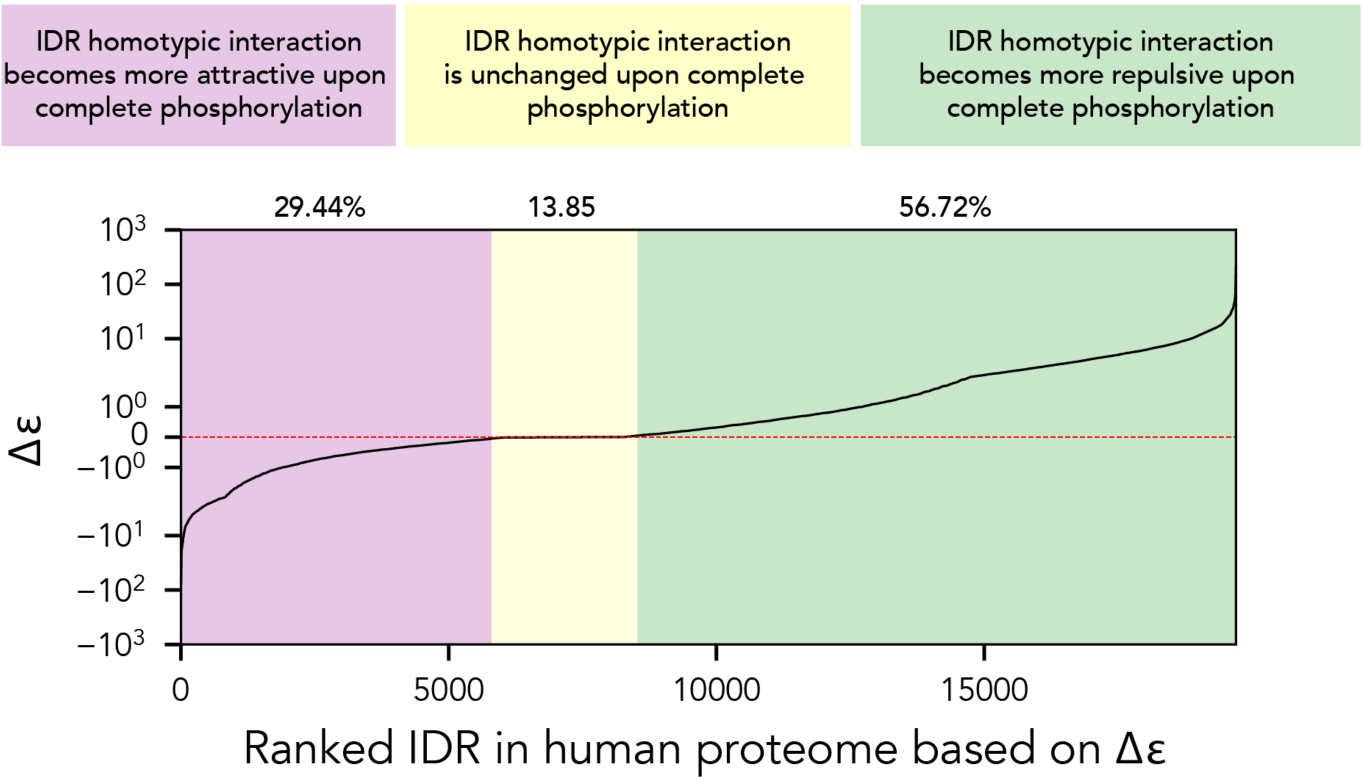


**Fig. S8 Changes in ϵ upon complete phosphorylation of an IDR**. We identified all IDRs in the human proteome that possess one or more experimentally-identified phosphosite (19,703 sequences). For each sequence, we computed the homotypic ϵ value prior to and upon complete phosphorylation of all phosphosites. The figure reports the ranked change in ϵ values upon phosphorylation; negative values imply ϵ becomes more attractive upon phosphorylation, while positive values imply ϵ becomes more attractive. We find many (thousands) of sequences where homotypic interaction is enhanced or suppressed upon phosphorylation. While this result suggests that the phosphorylation state can inherently tune self-interaction in IDRs, we make no claims as to whether homotypic intermolecular interaction has any physiological relevance. Instead, our interpretation of this result is that intermolecular interaction, *in general,* can be modulated by phosphorylation. Moreover, we expect that if a partner is known, the likely direction of that change (if driven by chemical specificity) can be predicted. For example, among our analysis's top hits is the IDR from Zona Occludens 1 (ZO1, UniProt ID Q07157). Our analysis predicts $ZO1^{798\text{-}1650}$ (the 853 residue C-terminal IDR) will have a dramatic reduction in homotypic interaction upon phosphorylation (Δϵ +73.2), a result strongly supported by extant data (*44*).

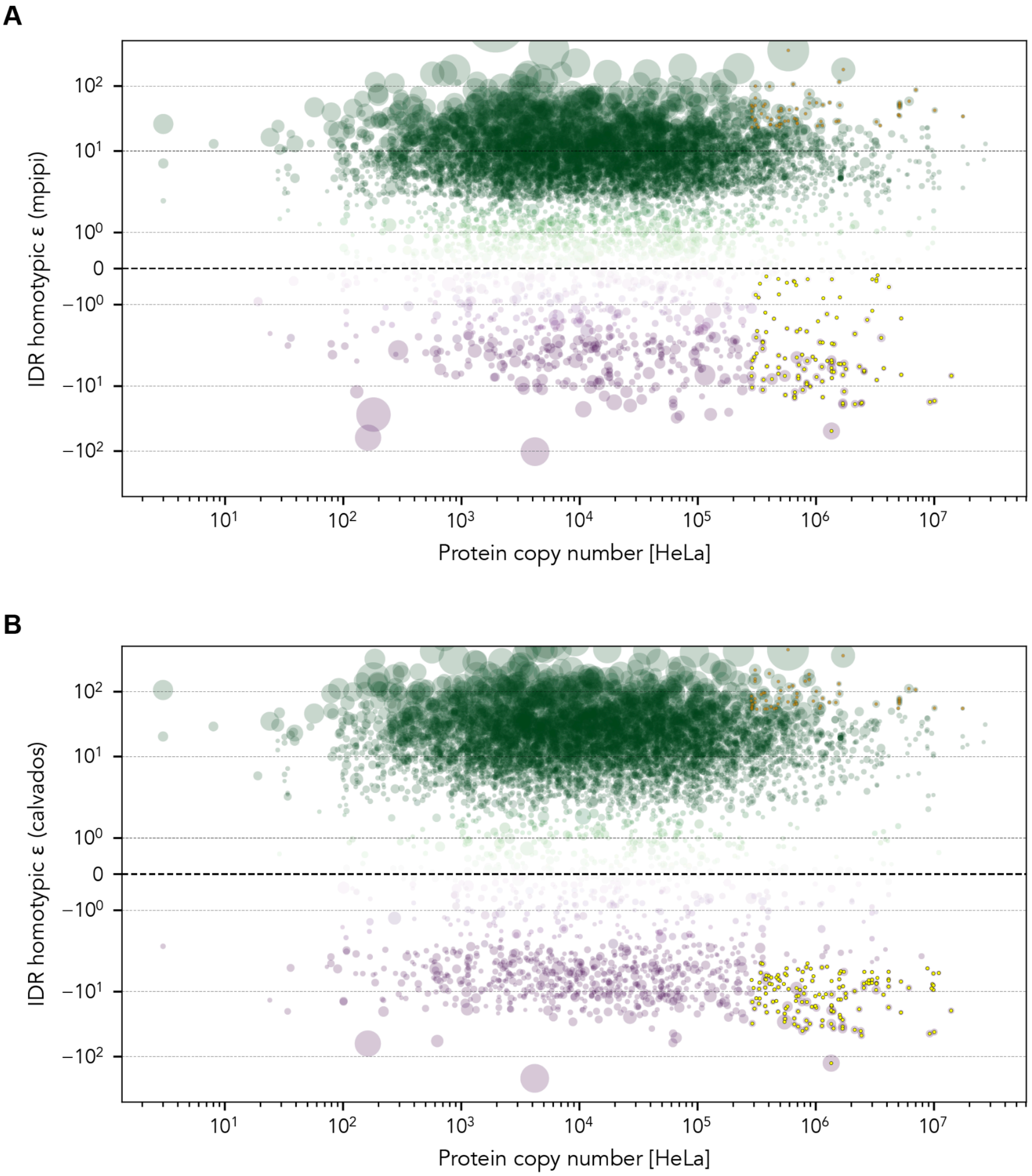


**Fig. S9 Comparison of homotypic ϵ vs. proteins copy number. (A, B)** Assessment of homotypic ϵ value for all IDRs in the human proteome where copy number data are available from Hein et al., giving a total of 8,088 IDRs (*15*). Marker size reports on IDR length, while the color reports on the ϵ value. The highly abundant points (present at over 284,000 copies per cell - in the top 10% of proteins) and highly attractive or repulsive (in the top 8% in terms of ϵ attraction or ϵ repulsion) are highlighted in yellow and orange.

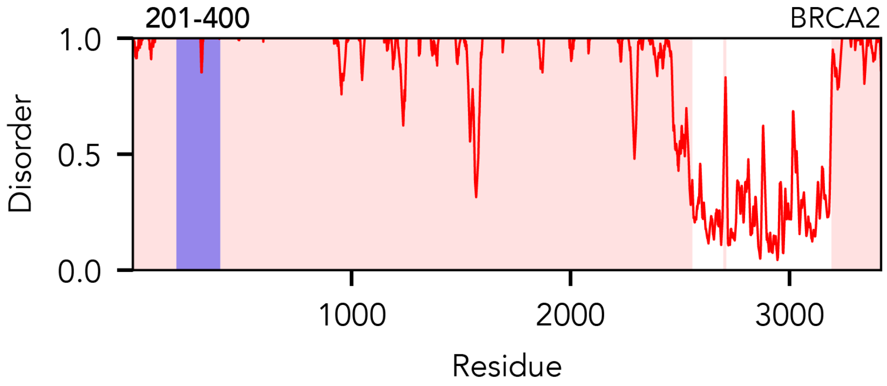


**Fig. S10 BRCA2 disorder profile**. BRCA2 is used as an example sequence for domain decomposition in Fig. 4 - disorder profile for the full-length protein along with the specific subregion of interest (200-400) is shown here.

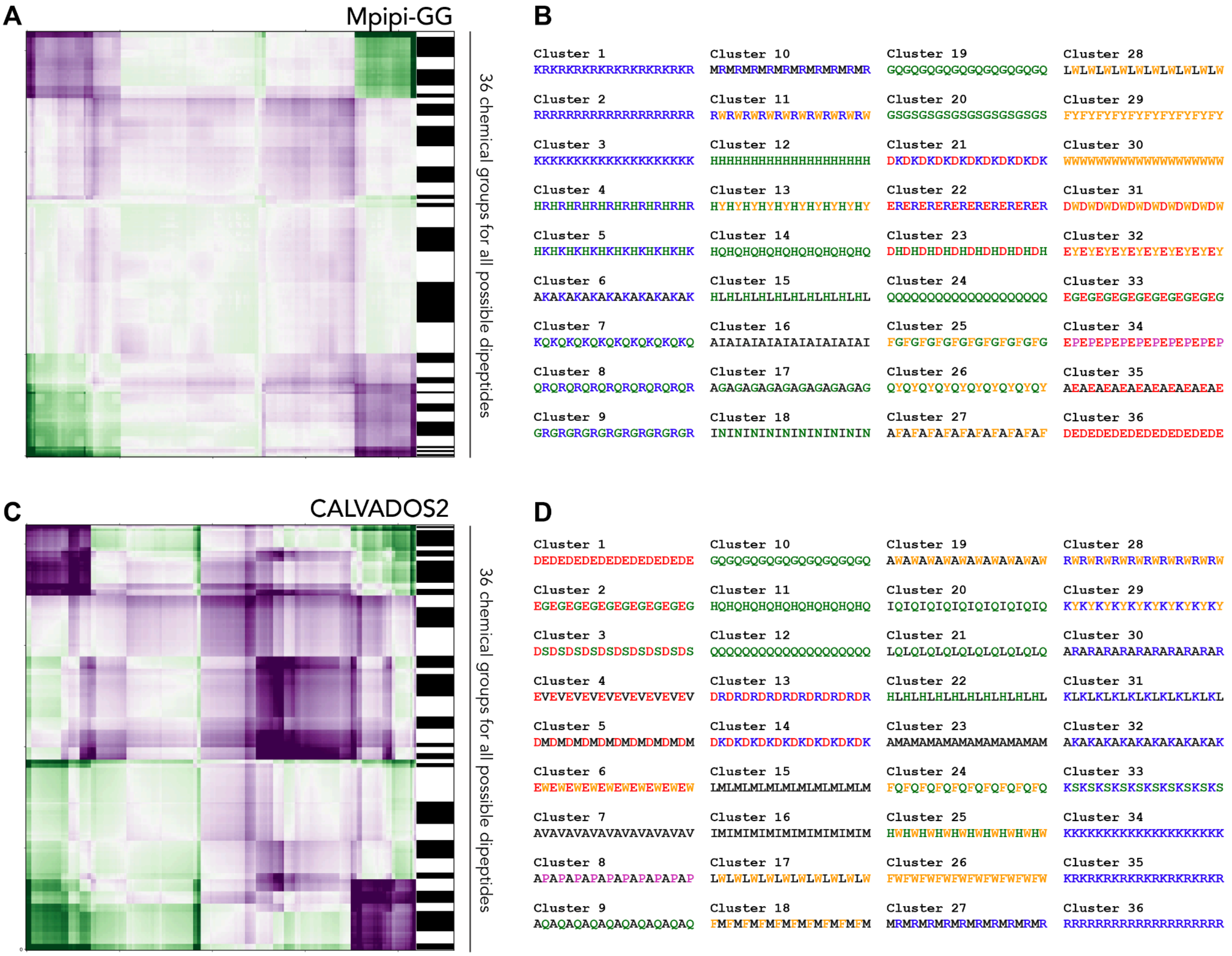


**Fig. S11 Chemical fingerprint clustering.** (**A,C**). All possible dipeptide repeats (10 repeats, 20 residues) had their heterotypic and homotypic ϵ values calculated. The resulting all vs. all matrix was hierarchically clustered into 36 groups with chemically similar dipeptides. (**B,D**) Representative dipeptides from each of the chemical groups. These 36 dipeptides provide a chemically orthogonal basis set, such that for a given IDR of interest, the heterotypic ϵ values between a guest IDR and all 36 fingerprints can be calculated to position the IDR of interest in 36-dimensions space. IDRs can then be compared in terms of their distance apart in this 36-dimensional space. This provides a general approach for measuring IDR similarity in chemical space.

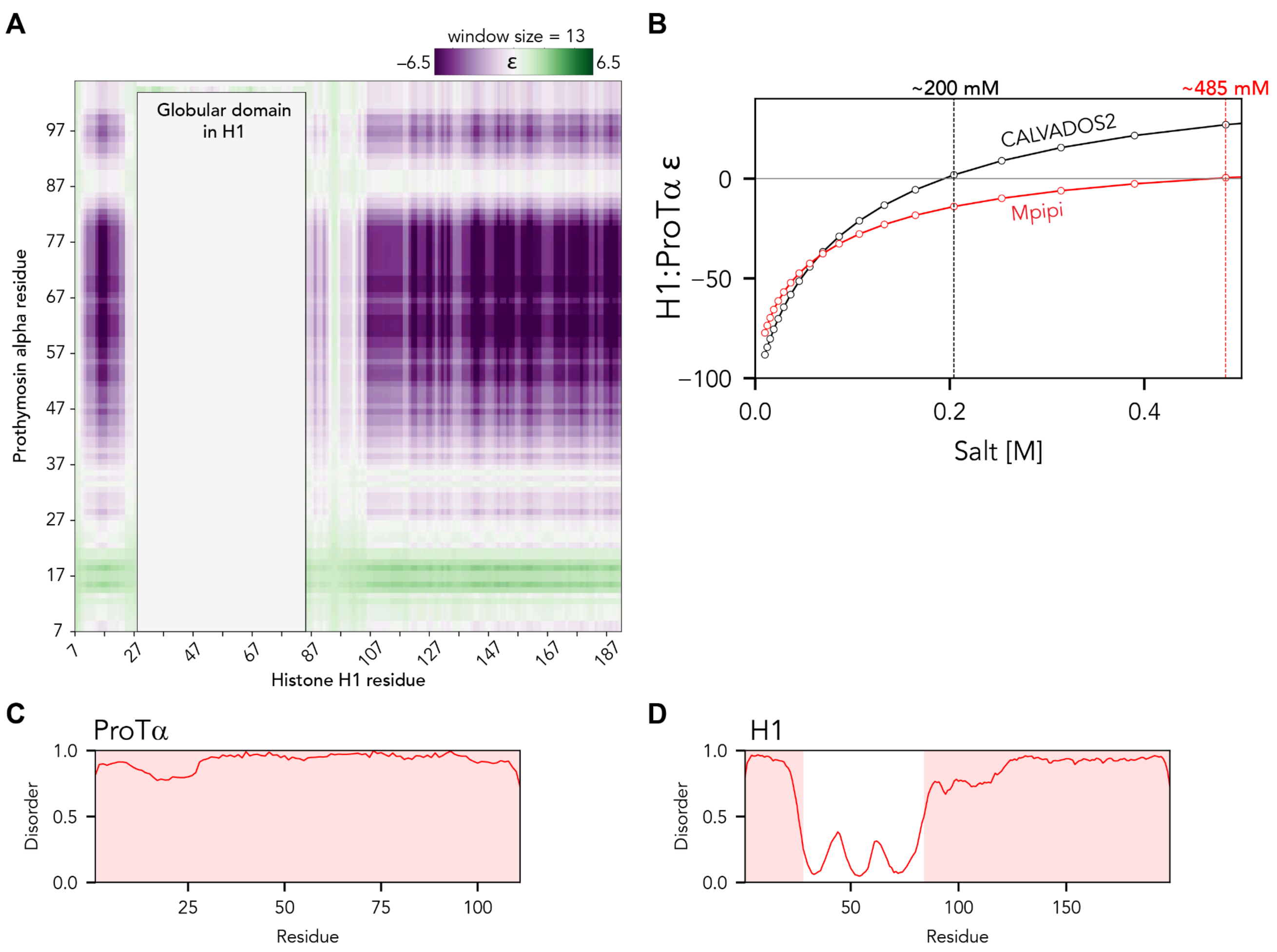


**Fig. S12 Histone H1:ProTα interaction. (A)** Heterotypic H1:ProTα intermap, showing which regions of both proteins interact. Summed intermaps are used to project binding regions onto a single axis in **Fig. 5**. This intermap was calculated using a sliding window of size 13. **(B)** Salt-dependence of H1:ProTα ϵ value. The value of 200 mM NaCl obtained by CALVADOS2 is in good agreement with the inferred critical salt concentration obtained by Galvanetto & Ivanović et al. (*45*). While the Mpipi-GG salt dependence is in less good quantitative agreement, both models predict a substantial change in mean-field intermolecular interaction between 0 and 150 mM NaCl, in excellent agreement with experimental work. (**C,D**) Predicted Prothymosyn α and Histone H1 disorder profiles.

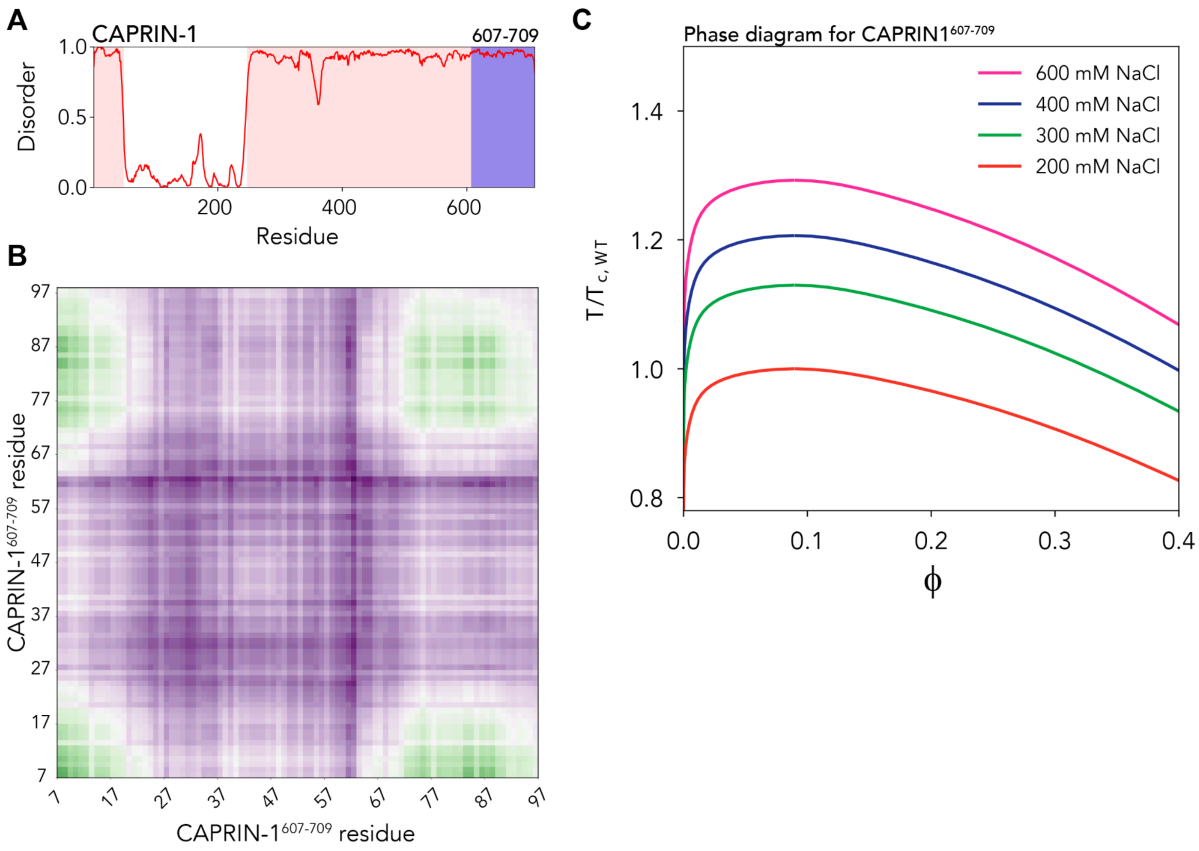


**Fig. S13 CAPRIN-1$^{607-709}$ interaction and phase behavior. (A)** Disorder profile for full-length CAPRIN-1, with the C-terminal IDR (residues 607-709) highlighted. **(B)** Homotypic intermap for CAPRIN-1$^{607-709}$. This intermap was calculated using a sliding window of size 13. (C) Salt dependence of CAPRIN-1 phase diagrams mirrors results obtained experimentally by Kim *et al.* - note here that increasing salt enhances the driving force for phase separation by suppressing electrostatic repulsion (*46*).

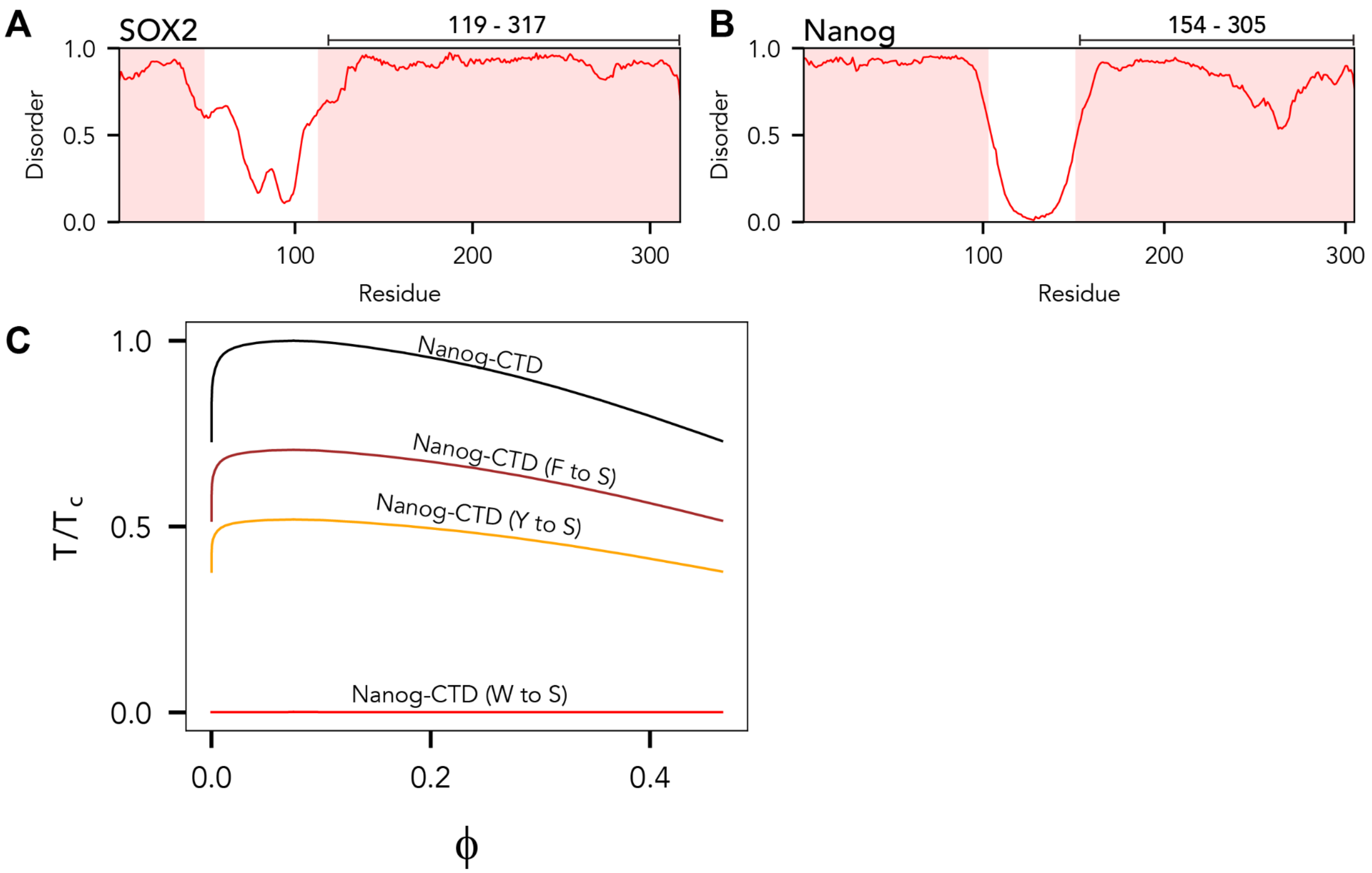


**Fig. S14 SOX2 and Nanog domain structure and phase behavior (A, B)** Disorder profiles for SOX2 and Nanog, with the two regions of interest highlighted. **(C)** Predicted phase diagrams for Nanog CTD with different aromatic mutant variants. These results recapitulate experimental trends reported previously (*47*).

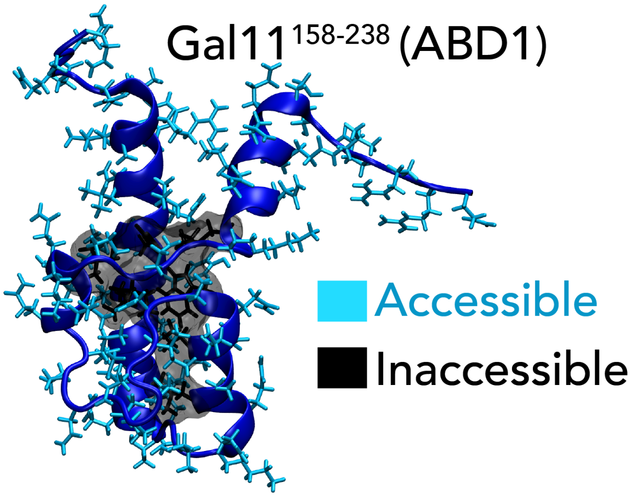


**Fig. S15** Gal11 activation-domain binding domain 1 (ABD1) with accessible and inaccessible residues highlighted. Intermolecular interactions were only calculated by a tile sequence and the accessible residues.

**sv1**
EKEKEKEKEK EKEKEKEKEK EKEKEKEKEK EKEKEKEKEK EKEKEKEKEK

**sv2**
EEEKKKEEEK KKEEEKKKEE EKKKEEEKKK EEEKKKEEEK KKEEEKKKEK

**sv3**
KEKKKEKKEE KKEEKEKEKE KEEKKKEEKE KEKEKKKEEK EKEEKKEEEE

**sv4**
KEKEEKEKKK EEEEKEKKKK EEKEKEKEKE EKKEEKKKKE EKEEKEKEKE

**sv5**
KEKEKKEEKE KKEEEKKEKE KEKKKEEKKK EEKEEKKEEK KKEEKEEEKE

**sv6**
EEEKKEKKEE KEEKKEKKEK EEEKKKEKEE KKEEEKKKEK EEEEKKKKEK

**sv7**
EEEEKKKKEE EEKKKKEEEE KKKKEEEEKK KKEEEEKKKK EEEEKKKKEK

**sv8**
KKKKEEEEKK KKEEEEKKKK EEEEKKKKEE EEKKKKEEEE KKKKEEEEKE

**sv9**
EEKKEEEKEK EKEEEEEKKE KKEKKEKKKE EKEKEKKKEK KKKEKEEEKE

**sv10**
EKKKKKKEEK KKEEEEEKKK EEEKKKEKKE EKEKEEKEKK EKKEEKEEEE

**sv11**
EKEKKKKKEE EKKEKEEEEK EEEEKKKKKE KEEEKEEKKE EKEKKKEEKK

**sv12**
EKKEEEEEEK EKKEEEEKEK EKKEKEEKEK KEKKKEKKEE EKEKKKKEKK

**sv13**
KEKKKEKEKK EKKKEEEKKK EEEKEKKKEE KKEKKEKKEE EEEEEKEEKE

**sv14**
EKKEKEEKEE EEKKKKKEEK EKKEKKKKEK KKKKEEEEEE KEEKEKEKEE

**sv15**
KKEKKEKKKE KKEKKEEEKE KEKKEKKKKE KEKKEEEEEE EEKEEKKEEE

**sv16**
EKEKEEKKKE EKKKKEKKEK EEKKEKEKEK KEEEEEEEEE KEKKEKKKKE

**sv17**
EKEKKKKKKE KEKKKKEKEK KEKKEKEEEK EEKEKEKKEE KKEEEEEEEE

**sv18**
KEEKKEEEEE EEKEEKKKKK EKKKEKKEEE KKKEEKKKEE EEEEKKKKEK

**sv19**
EEEEEKKKKK EEEEEKKKKK EEEEEKKKKK EEEEEKKKKK EEEEEKKKKK

**sv20**
EEKEEEEEEK EEEKEEKKEE EKEKKEKKEK EEKKEKKKKK KKKKKKKEEE

**sv21**
EEEEEEEEEK EKKKKKEKEE KKKKKKEKKE KKKKEKKEEE EEEKEEEKKK

**sv22**
KEEEEKEEKE EKKKKEKEEK EKKKKKKKKK KKKEKKEEEE EEEEKEKEEE

**sv23**
EEEEEKEEEE EEEEEEEKEE KEKKKKKKEK KKKKKKEKEK KKKEKKEEKK

**sv24**
EEEEKEEEEE KEEEEEEEEE EEEKKKEEKK KKKEKKKKKK KEKKKKKKKK

**sv25**
EEEEEEEEEE EKEEEEKEEK EEKEKKKKKK KKKKKKKKKK KKEEKKEEKE

**sv26**
KEEEEEEEKE EKEEEEEEEE EKEEEEKEEK KKKKKKKKKK KKKKKKKKKE

**sv27**
KKEKKKEKKE EEEEEEEEEE EEEEEEEEEK EEKKKKKKKK KKKKKKKEKK

**sv28**
EKKKKKKKKK KKKKKKKKKK KKEEEEEEEE EEEEEEEEEE KKEEEEEEKEK

**sv29**
KEEEEKEEEE EEEEEEEEEE EEEEEEEKKK KKKKKKKKKK KKKKKKKKKK

**sv30**
EEEEEEEEEE EEEEEEEEEE EEEEEKKKKK KKKKKKKKKK KKKKKKKKKK

**Fig. S16** Das-Pappu sequences, as reported previously (*8*).

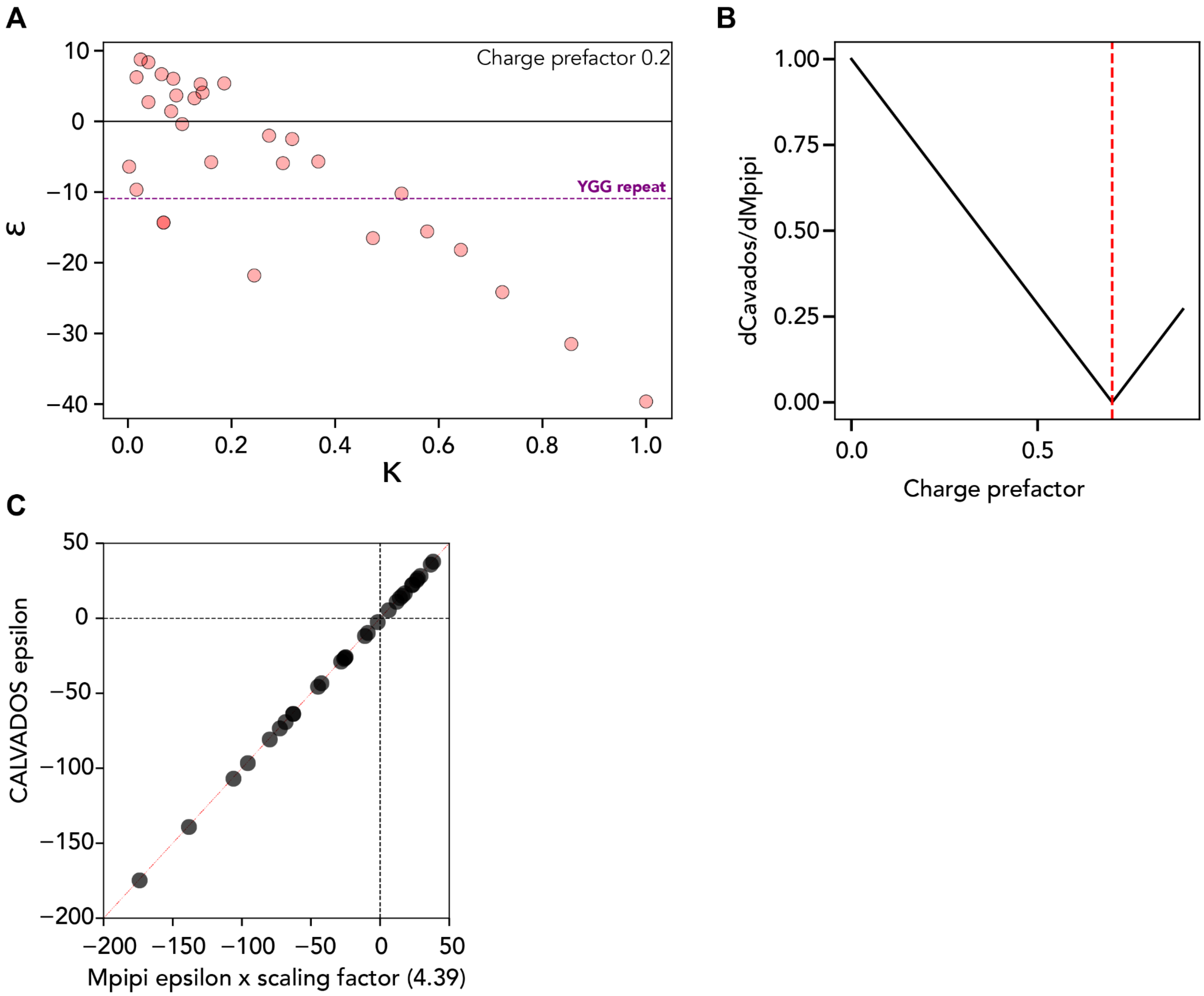


**Fig. S17 FINCHES calibration. (A)** Mpipi-GG ϵ values for Das-Pappu sequences under 0 mM salt using a charge prefactor of 0.2. The YGG repeat sequence is used as a point comparison. **(B)** Comparison between Mpipi-GG and CALVADOS2 predictions for Das-Pappu sequences with different charge prefactors for CALVADOS2. We reached the best agreement when the CALVADOS2 prefactor is set to 0.7. **(C)** With a prefactor to correct for the difference in magnitude for CALVADOS2 and Mpipi-GG, under charge prefactors of 0.2 and 0.7 Mpipi-GG and CALVADOS2 show 1:1 correlation in terms of ϵ with respect to Das-Pappu sequences.

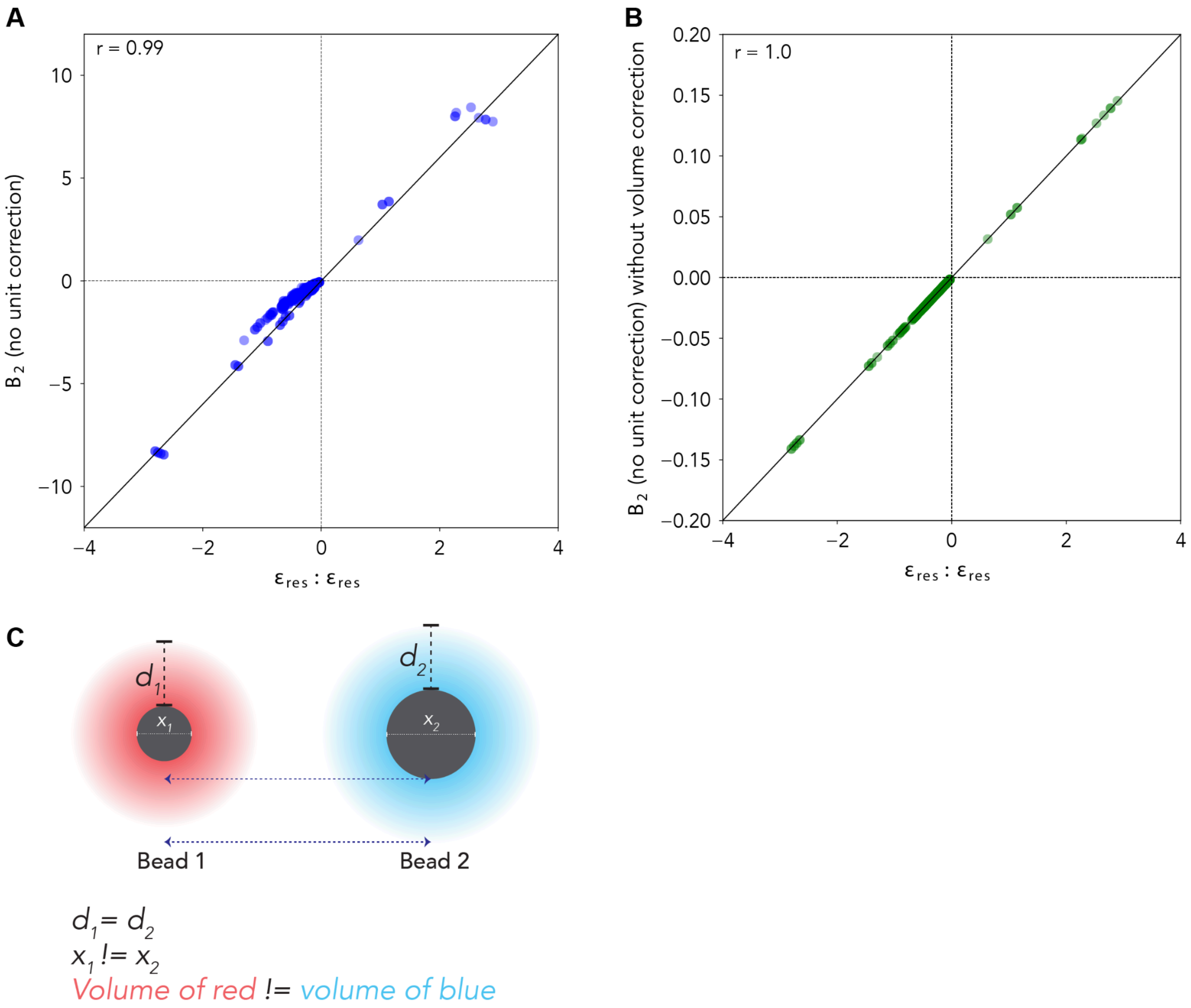


**Fig. S18 Comparison of $B_2$ vs. ϵ. (A)** Correlation between inter-bead analytical $B_2$ value (from the Mayer-f function) vs. ϵ. **(B)** Comparison of $B_2$ sans volume element correction vs. ϵ. **(C)** Schematic of beads with equivalent sized potentials ($d_1 = d_2$), but because $x_2 > x_1$ if a volume element correction were included, bead two would have a stronger intermolecular interaction than bead 1.

31. J. Abramson, J. Adler, J. Dunger, R. Evans, T. Green, A. Pritzel, O. Ronneberger, L. Willmore, A. J. Ballard, J. Bambrick, S. W. Bodenstein, D. A. Evans, C.-C. Hung, M. O’Neill, D. Reiman, K. Tunyasuvunakool, Z. Wu, A. Žemgulytė, E. Arvaniti, C. Beattie, O. Bertolli, A. Bridgland, A. Cherepanov, M. Congreve, A. I. Cowen-Rivers, A. Cowie, M. Figurnov, F. B. Fuchs, H. Gladman, R. Jain, Y. A. Khan, C. M. R. Low, K. Perlin, A. Potapenko, P. Savy, S. Singh, A. Stecula, A. Thillaisundaram, C. Tong, S.

Yakneen, E. D. Zhong, M. Zielinski, A. Žídek, V. Bapst, P. Kohli, M. Jaderberg, D. Hassabis, J. M. Jumper, Accurate structure prediction of biomolecular interactions with AlphaFold 3. *Nature*, doi: 10.1038/s41586-024-07487-w (2024).

46. T. H. Kim, B. J. Payliss, M. L. Nosella, I. T. W. Lee, Y. Toyama, J. D. Forman-Kay, L. E. Kay, Interaction hot spots for phase separation revealed by NMR studies of a CAPRIN1 condensed phase. *Proc. Natl. Acad.*

*Sci. U. S. A.* **118** (2021).

47. K.-J. Choi, M. D. Quan, C. Qi, J.-H. Lee, P. S. Tsoi, M. Zahabiyon, A. Bajic, L. Hu, B. V. V. Prasad, S.-C. J. Liao, W. Li, A. C. M. Ferreon, J. C. Ferreon, NANOG prion-like assembly mediates DNA bridging to facilitate chromatin reorganization and activation of pluripotency. *Nat. Cell Biol.* **24**, 737–747 (2022).
